# Synaptic plasticity in human thalamocortical assembloids

**DOI:** 10.1101/2024.02.01.578421

**Authors:** Mary H. Patton, Kristen T. Thomas, Ildar T. Bayazitov, Kyle D. Newman, Nathaniel B. Kurtz, Camenzind G. Robinson, Cody A. Ramirez, Alexandra J. Trevisan, Jay B. Bikoff, Samuel T. Peters, Shondra M. Pruett-Miller, Yanbo Jiang, Andrew B. Schild, Anjana Nityanandam, Stanislav S. Zakharenko

## Abstract

Synaptic plasticities, such as long-term potentiation (LTP) and depression (LTD), tune synaptic efficacy and are essential for learning and memory. Current studies of synaptic plasticity in humans are limited by a lack of adequate human models. Here, we modeled the thalamocortical system by fusing human induced pluripotent stem cell–derived thalamic and cortical organoids. Single-nucleus RNA-sequencing revealed that most cells in mature thalamic organoids were glutamatergic neurons. When fused to form thalamocortical assembloids, thalamic and cortical organoids formed reciprocal long-range axonal projections and reciprocal synapses detectable by light and electron microscopy, respectively. Using whole-cell patch-clamp electrophysiology and two-photon imaging, we characterized glutamatergic synaptic transmission. Thalamocortical and corticothalamic synapses displayed short-term plasticity analogous to that in animal models. LTP and LTD were reliably induced at both synapses; however, their mechanisms differed from those previously described in rodents. Thus, thalamocortical assembloids provide a model system for exploring synaptic plasticity in human circuits.

**Highlights:**

- Human thalamic organoids consist of mostly glutamatergic projection neurons.
- Thalamocortical assembloids form reciprocal glutamatergic synapses.
- Synapses are functional and undergo short-term plasticity resembling animal models.
- Long-term potentiation and depression reveal mechanisms distinct from rodents.

**eTOC:** Human organoids are often used to model diseases with synaptic pathology; however, few studies have examined synaptic function via single-cell or single-synapse recordings. Patton et al. fused human thalamic and cortical organoids into assembloids to examine synaptic transmission and short- and long-term synaptic plasticity in human thalamocortical and corticothalamic circuits.

**GRAPHICAL ABSTRACT:** 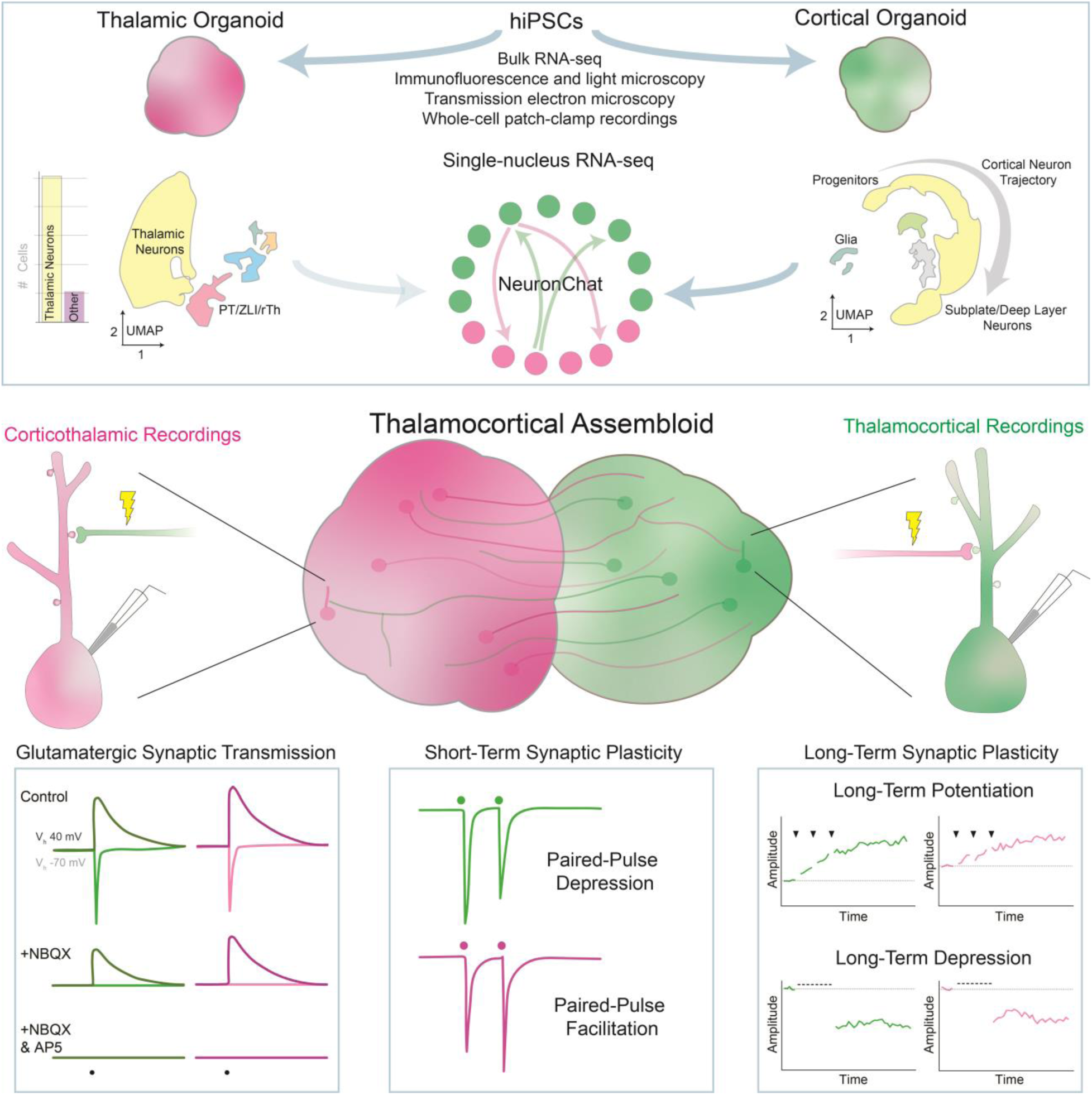

## INTRODUCTION

Synaptic transmission through the release of neurotransmitter and the subsequent activation of postsynaptic receptors is the fundamental mode of communication between neurons. Synaptic plasticity enhances or diminishes synaptic transmission in an activity-dependent manner in response to environmental cues. Although the chemical synapse is evolutionarily conserved,^1^ the molecular mechanisms underlying synaptic plasticity differ across species.^2–6^ Aberrant synaptic plasticity is well documented in animal models of autism, schizophrenia, and other psychiatric disorders.^7–10^ Ultimately, these human disorders would benefit from a human model system.

*Ex vivo* human model systems are currently limited to postmortem samples, which are often in a state of uncertain tissue quality, and biopsy samples, which are almost exclusively from individuals with epilepsy or brain tumors.^11^ The recent development of human induced pluripotent stem cell (hiPSC)-derived organoids provides a promising *in vitro* model system that is accessible to experimental manipulation and recapitulates human-specific features of neural development and function.^12–18^ Organoids are widely used to model synaptopathies (i.e., neurologic and psychiatric disorders associated with synaptic dysfunction).^19–25^ Specific properties of synaptic transmission and synaptic plasticity differ across brain regions and neuronal subtypes;^2,26–28^ however, few studies have examined synaptic transmission between defined pre- and postsynaptic cell populations in organoids.^29–31^ Moreover, synaptic plasticity within organoids has never been reported using whole-cell patch-clamp electrophysiology, the gold standard approach for studying synaptic physiology that has been validated over decades of research in animal models.

To build a functional neural circuit, human cortical organoids (hCOs) can be fused with human thalamic organoids (hThOs) to produce thalamocortical (TC) assembloids.^32–34^ The thalamus is a primary relay center for incoming sensory information that sends widespread, yet highly organized, projections to various cortical regions based on the sensory modality.^35–38^ The cortex, in turn, sends projections back to thalamic nuclei to integrate and update sensory, motor, and cognitive information.^39,40^ Collectively, synaptic transmission within the thalamo–cortico– thalamic circuit creates cognitive representations of the outside world based on diverse incoming sensory inputs and provides the foundation for dynamic executive functioning.^41–44^ Experience-dependent synaptic plasticity within this system is critical to learning and underlies the expression of sensorimotor behaviors, attention, and perceptual and working memory.^44–49^

Here we developed a human assembloid system containing functional glutamatergic TC and corticothalamic (CT) synaptic connections that undergo short- and long-term synaptic plasticity. We then used this system to explore the molecular mechanisms underlying synaptic plasticity in human neurons.

## RESULTS

### hThOs contain functional glutamatergic thalamic neurons

To build optimal hThOs, we generated a reporter line from an hiPSC line derived from a neurotypical male subject with normal karyotype (See **Figure S1** for reporter line validation). Specifically, the *tdTomato-*coding sequence was inserted into the endogenous *TCF7L2* locus (**Figure 1A**). We then modified a previously reported protocol^50^ to increase the efficiency of thalamic neuron generation. After differentiation into hThOs, we performed bulk RNA-sequencing and then VoxHunt deconvolution analysis of the RNA-seq data. High *tdTomato* RNA levels identified hThOs with high representation of diencephalon, the developmental structure that gives rise to the thalamus, and low representation from contaminating structures, e.g., the pallium and midbrain (**Figure 1A**). Notably, in hThOs generated from five independent hiPSC lines, *TCF7L2* positively predicted the expression of the thalamic neural precursor genes *OLIG3* and *OTX2* and the thalamic neuron genes *GBX2* and *LHX9* (**Figure S2**). Visual assessment was sufficient to identify hThOs with high *tdTomato*, as differential expression analysis comparing hThOs with high TCF7L2-tdTomato fluorescence to hCOs revealed significant enrichment of thalamic markers in the hThOs (**Figure S3A** and **S3B**). All subsequent experiments were performed with TCF7L2-tdTomato^+^ hThOs.

**Figure 1.**
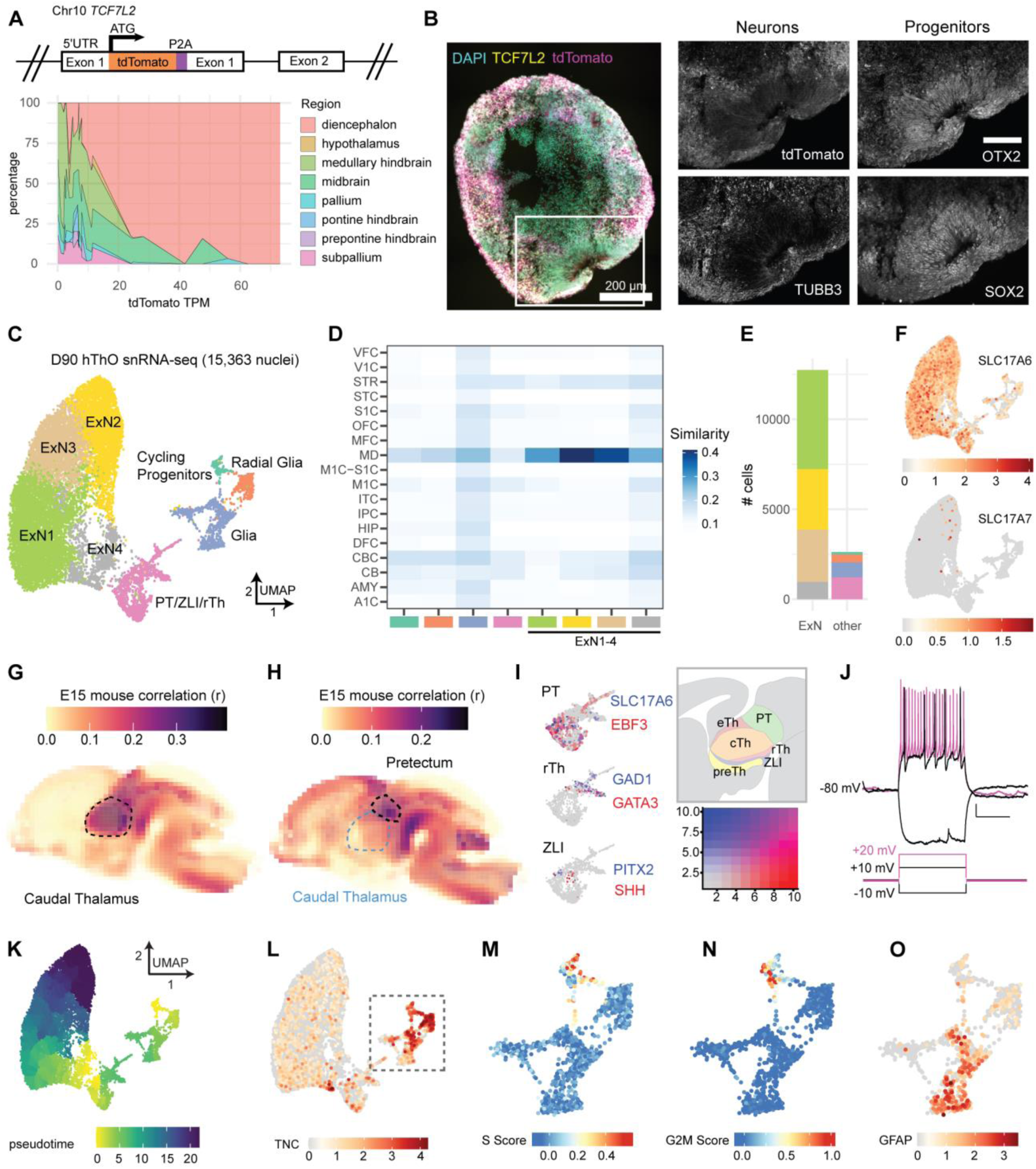
hThOs contain functional glutamatergic thalamic neurons. (**A**) Reporter cell line validation for hThOs. Top: Schematic of *TCF7L2* exon 1 in the TP-190a-TCF7L2-tdTomato reporter line, which was used to generate all hThOs, except where indicated in Figure S2. Bottom: VoxHunt deconvolution analysis of bulk RNA-seq data from D69-D70 hThOs using E13 mouse brain data from the *Allen Brain Atlas* as a reference. (**B**) Immunofluorescence images of TCF7L2, TUBB3, OTX2, and SOX2 labeling in D60 hThOs. Nuclei are indicated by DAPI (cyan). TCF7L2-tdTomato fluorescence is indicated in magenta. Images were acquired from serial sections of the same organoid. Scale bars: 200 µm (whole section), 100 µm (insets). (**C**) UMAP plot with cluster annotations indicated by color. (**D**) VoxHunt analysis by snRNA-seq cell cluster. Excitatory neuron (ExN) clusters exhibit the highest correlations with *BrainSpan* samples from human mediodorsal nucleus of the thalamus (MD), aged 13–24 pcw. Cluster annotations are indicated by color on the x-axis. (**E**) Bar plot showing the number of nuclei per cell cluster, with clusters indicated by fill color. (**F**) UMAP plots of glutamatergic markers *SLC17A6* (*VGLUT2*) and *SLC17A7* (*VGLUT1*). Color indicates normalized transcript level. (**G**) VoxHunt correlation analysis mapping clusters ExN1-4 onto the E15 mouse brain. (**H**) VoxHunt correlation analysis mapping the PT/ZLI/rTh cluster onto the E15 mouse brain. (**I**) UMAP plots of the PT/ZLI/rTh cluster, demonstrating the expression of markers associated with the PT, ZLI, and rTh. Transcript information is indicated by color. The relative locations of these structures within the developing diencephalon are shown in the schematic. (**J**) Example traces showing voltage and AP responses to current injections in a cell recorded from an hThO. (**K**) Pseudotime ordering of cells within the hThOs. (**L**) UMAP plot of the neural progenitor marker *TNC*. Color indicates the normalized transcript level. (**M-N**) UMAP plots of cell cycle analysis results for the Cycling Progenitor, Radial Glia, and Glia cell clusters. Color indicates S Score (**M**) or G2M Score (**N**). (**O**) UMAP plot of the astrocyte marker *GFAP* in the Cycling Progenitor, Radial Glia, and Glia cell clusters. Color indicates the normalized transcript level. Data in (**C-I**) and (**K-O**) were produced by snRNA-seq analysis of 15,363 nuclei from D90 hThOs. See **Figures S1-S3** for additional data validating hiPSC lines and hThOs. See **Figure S4** for additional data related to electrophysiological properties and synapses in hThOs.

At 60 days after the start of differentiation (D60), hThOs contained OTX2^+^ and SOX2^+^ neural progenitor domains surrounded by TUBB3^+^ neurons (**Figure 1B**). By D92, most cells expressed markers consistent with glutamatergic neurons of the developing thalamus, specifically LHX2, FOXP2, and GBX2; GABA immunoreactivity was observed in only a small subset of cells (**Figure S3C**). At D70–D90 in culture, synaptic gene expression significantly increased, and precursor and mitotic gene expression decreased in hThOs (**Figure S3D and S3E**).

Whereas previous protocols generated hThOs containing ∼25-50% glutamatergic neurons,^32,33^ single-nucleus RNA-sequencing (snRNA-seq) analysis of our D90 hThOs revealed that the majority (∼85%) of cells were glutamatergic neurons found primarily in Excitatory Neuron 1-4 (ExN1-4) clusters (**Figure 1C-1F**). Applying VoxHunt mapping to *BrainSpan* and *Allen Brain Atlas* references, we found that clusters ExN1-4 mapped to the human mediodorsal thalamus (**Figure 1D**) and embryonic day (E) 15 mouse caudal thalamus (**Figure 1G**), the diencephalon structure that produces the glutamatergic neurons of the mature thalamus. Consistent with thalamic neurons, ExN1-4 neurons primarily expressed *SLC17A6* (or *VGLUT2*); *SLC17A7* (or *VGLUT1*) was sparsely expressed (**Figure 1F**). Neurons in clusters ExN1-4 expressed additional thalamic markers of interest, including *GBX2*, *SHOX2*, *FOXP2*, *CADM1*, and *NTNG1* (**Figure S3F**). In line with reports from the mouse thalamus,^51^ a subset of cells also expressed *SOX2*, a marker typically associated with precursors rather than mature neurons (**Figure S3F**). We also identified one cluster that mapped to E15 mouse pretectum (PT) (**Figure 1H**), as well as cells expressing markers of the thalamic organizer zona limitans intrathalamica (ZLI) and rostral thalamus (rTh) (**Figure 1I**). Like the caudal thalamus, these structures arise from the diencephalon during early development. A small subset of *SLC17A6^+^* glutamatergic neurons and all *GAD1*^+^ GABAergic neurons within the hThOs were found in this PT/ZLI/rTh cluster.

To verify that the predominant cells in hThOs were functional putative neurons, we used whole-cell patch-clamp electrophysiology to investigate the membrane properties of individual cells. Action potentials (APs) were evoked in response to depolarizing current injections (**Figure 1J**). The resting membrane potential, membrane capacitance, input resistance, and AP properties measured in hThO cells were consistent with neurons (**Figure S4A-S4H**). Transmission electron microscopy (TEM) also revealed numerous asymmetric and symmetric synapses in hThOs (**Figure S4Q** and **S4R**). A subset of presynaptic terminals contained dense core vesicles (**Figure S4S**).

Finally, we examined the non-neuronal cell populations in our hThOs, which together constituted ∼15% of cells. Pseudotime analysis (**Figure 1K**) and cell cycle analysis (**Figure 1L-1N**) revealed a small cluster of precursor cells undergoing mitosis (i.e., Cycling Progenitors cluster). *TNC^+^* progenitors not undergoing mitosis were labeled radial glia. Most of the remaining cells were *GFAP^+^* astrocytes within the Glia cluster (**Figure 1O**). Pseudotime analysis also revealed differences in glutamatergic neuron maturity, with the ExN4 cluster containing the least mature thalamic neurons, and the ExN2 cluster containing the most mature thalamic neurons (**Figure 1K**). We conclude that our hThO protocol generated functional thalamic neurons with high efficiency and that those neurons formed synaptic connections.

### TC assembloids form functional glutamatergic TC and CT synapses

To form TC assembloids, we also required optimal hCOs. We generated an isogenic hiPSC reporter line by inserting the *tdTomato-*coding sequence into the endogenous *SLC17A7* (*VGLUT1*) locus (**Figure 2A**) (See **Figure S1** for reporter line validation). Using this reporter line, we generated hCOs via a previously reported protocol.^52^ VoxHunt deconvolution analysis of bulk RNA-seq data demonstrated that VGLUT1-tdTomato^+^ hCOs exhibited high representation of the pallium, the developmental structure that gives rise to the neocortex, and low representation of contaminating structures, e.g., the subpallium and diencephalon (**Figure 2A**). High expression of *tdTomato* RNA also predicted high expression of cortical markers relative to hThOs (**Figure S3A** and **S3B**) and VGLUT1-tdTomato^−^ hCOs (**Figure S5A**). All subsequent experiments were performed with VGLUT1-tdTomato^+^ hCOs.

**Figure 2.**
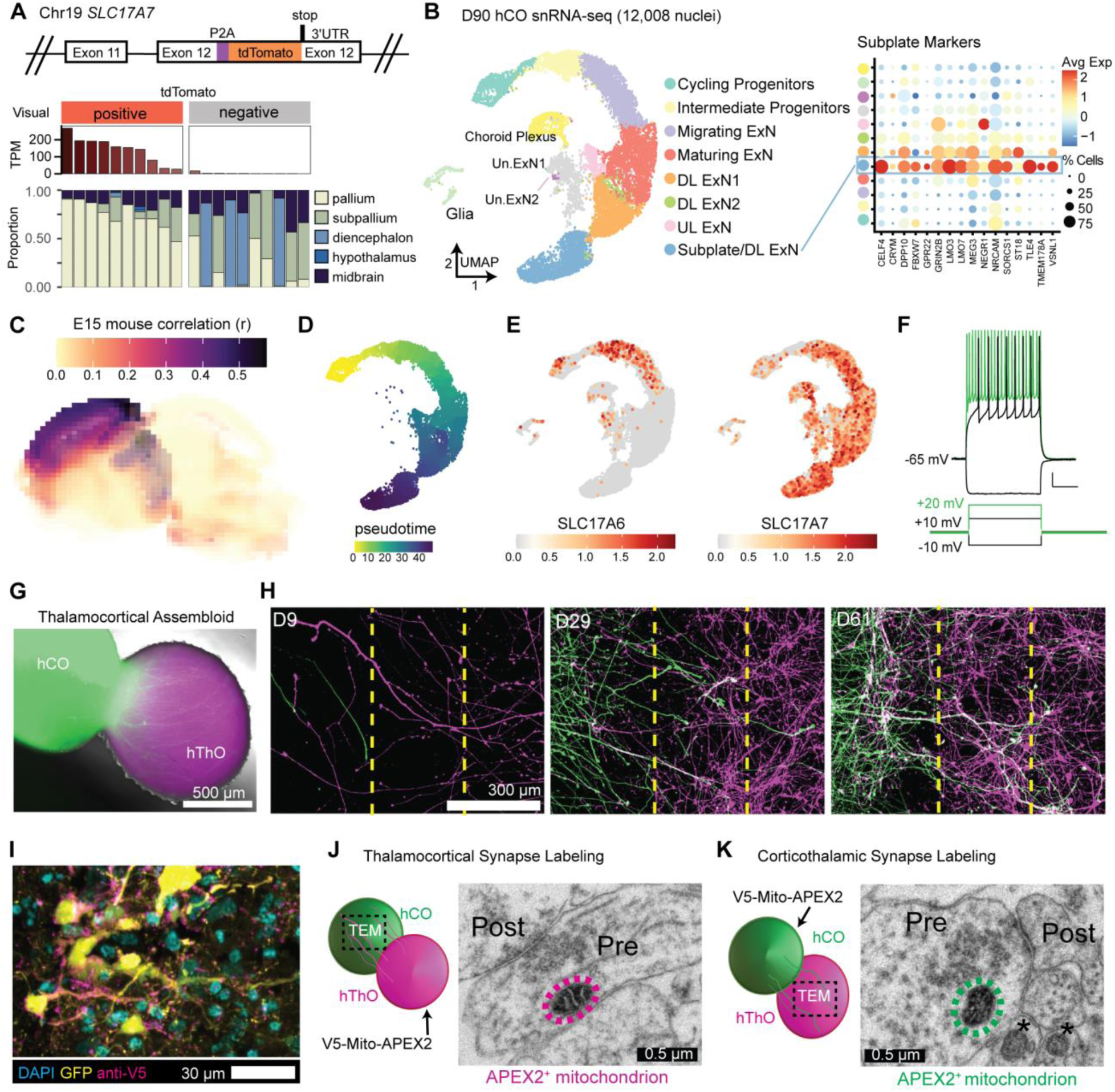
Fusing hThOs and hCOs produces assembloids that form reciprocal synapses. (**A**) Reporter line validation for hCOs. Top: Schematic of *SLC17A7* (*VGLUT1*) exon 12 in the TP-190a-VGLUT1-tdTomato reporter line, which was used to generate all hCOs. Bottom: VoxHunt deconvolution analysis of bulk RNA-seq data from D70 hCOs using E13 mouse brain data from the *Allen Brain Atlas* as a reference. Organoids were visually categorized as positive or negative for tdTomato fluorescence prior to sequencing. The *tdTomato* RNA level for each sample is indicated in TPM (transcripts per million). Each stacked bar indicates one bulk RNA-seq sample derived from 2-3 pooled organoids. (**B**) The snRNA-seq analysis of hCOs. Left: UMAP plot with cluster annotations. ExN: excitatory neuron, DL: deep layer, UL: upper layer, Un.: unknown. Right: Dot plot showing subplate marker expression by cluster. Avg Exp: normalized average expression, % Cells: percentage of cells expressing a marker within a cluster. (**C**) VoxHunt analysis mapping hCOs (all clusters) onto the E15 mouse brain. (**D**) Pseudotime analysis of the neural cell trajectory (Cycling Progenitors to UL ExNs, DL ExNs, and Subplate/DL ExNs) from hCOs. (**E**) UMAP plots of glutamatergic markers *SLC17A6* (*VGLUT2*) and *SLC17A7* (*VGLUT1*). Color indicates normalized transcript level. (**F**) Traces showing the voltage and AP responses in a cell recorded from an hCO. (**G**) Fluorescence and bright field image of a TC assembloid at 5 days postfusion (dpf). (**H**) Representative fluorescence images for 2-dimensional fusion assay. Thalamic neurons (magenta, right chamber) and cortical neurons (green, left chamber) extend processes from their respective chambers, across the barrier region (dashed yellow lines), and into the opposite chamber starting at D9. Elaborate processes extending from the opposite sides can be seen in both halves by D61. (**I**) Fluorescence image of an hCO co-transduced with hSyn-GFP and hSyn-V5-Mito-APEX2 lentiviruses. (**J**) Schematic and TEM image of an APEX2^+^ mitochondrion (circled in magenta) in a TC synapse. Pre: presynaptic compartment, post: postsynaptic compartment. (**K**) Schematic and TEM image of an APEX2^+^ mitochondrion (circled in green) in a CT synapse. APEX2^−^ mitochondria are indicated by asterisks (*). Scale bars (**F**): 10 mV, 2.5 s. Data in (**B-E**) were produced by snRNA-seq analysis of 12,008 nuclei from D90 hThOs. See **Figures S1 and S5** for additional data validating the TP-190a-VGLUT1-tdTomato reporter line and hCOs, respectively. See **Figure S4** for additional data related to electrophysiological properties and synapses in hCOs. See **Figure S6** for NeuronChat analysis.

The snRNA-seq analysis of D90 hCOs revealed that most cells (∼85%) fell within a neuronal developmental trajectory, beginning with neural precursors and ending with differentiated neurons that expressed markers of upper-layer (UL) or deep-layer (DL) excitatory cortical neurons (UL ExNs and DL ExNs, respectively) (**Figures 2B** and **S5**). Using VoxHunt mapping to *BrainSpan* and *Allen Brain Atlas* references, we found that clusters within this trajectory mapped to human neocortical structures (**Figure S5B**) and E15 mouse neocortex (**Figure 2C**). Consistent with previous reports in rodent models,^53^ *SLC17A6* was expressed in intermediate progenitors, but expression declined with neuronal maturation (**Figure 2E**). Mature neurons in hCOs exclusively expressed *SLC17A7* (**Figure 2F**). Remaining cells (∼15%) included unidentified glutamatergic neurons (found in the Un.ExN1 and Un.ExN2 clusters), glia, and cells resembling those in choroid plexus (**Figure S5I** and **S5J**). Overall, glutamatergic neurons constituted ∼78% of cells in hCOs.

During early development, thalamic neurons first form synapses within the cortical subplate before transitioning to cortical Layer IV.^54^ Conversely, subplate neurons project to several thalamic nuclei.^55^ Thus, cortical subplate neurons are critical to TC and CT circuitry development. We identified a cluster in hCOs (Subplate/DL ExN) that was enriched for markers of the cortical subplate^56^ (**Figure 2B**) and contained the most mature neurons based on pseudotime analysis (**Figure 2D**), which was in line with this cluster containing subplate-like neurons.^57^ We then applied NeuronChat^58^ to our snRNA-seq data to determine the likelihood of neuronal communication between cell clusters in the hThOs and hCOs. We found that the hThO ExN clusters, which contain glutamatergic thalamic neurons, exhibited the highest probability of TC communication with cells in the Cycling Progenitor and Subplate/DL ExN clusters of the hCOs (**Figure S6A**). Conversely, Subplate/DL ExNs exhibited a higher probability of CT communication with hThO ExN clusters than exhibited by other hCO clusters (**Figure S6B**). Interactions between thalamic axons and cortical progenitors during mouse development are well-documented but are not driven by synaptic connections.^59–61^ Further analysis revealed that hThO ExNs exhibited a high probability of TC communication with both hCO Cycling Progenitors and Subplate DL/ExNs clusters by NRXN signaling (**Figure S6C**). However, hThO ExNs exhibited a higher probability of glutamatergic communication with Subplate DL/ExNs than hCO Cycling Progenitors (**Figure S6D**). Together, these analyses suggest that hCOs contain neurons which might be capable of forming glutamatergic TC and CT synapses.

Next, we investigated the firing properties of hCO cells by using whole-cell patch-clamp electrophysiological recordings. Delivering depolarizing current injections to hCO cells evoked AP firing (**Figures 2F** and **S4L-S4P**). These cells displayed typical neuronal properties (**Figure S4I-S4P**), consistent with previous reports of hCOs.^62–64^ TEM revealed numerous asymmetric and symmetric synapses in hCOs (**Figure S4Q** and **S4R**). A subset of presynaptic terminals contained dense core vesicles (**Figure S4S**).

We then fused hThOs with hCOs to form TC assembloids. The hCOs were transduced with hSyn-GFP lentivirus prior to fusion, so each organoid could be identified within the assembloid (**Figure 2G**). GFP^+^ axons from the hCO were detectable within the hThO within 5 days postfusion (dpf). Furthermore, 2-dimensional fusion assays confirmed that hThOs and hCOs sent reciprocal axonal projections (**Figure 2H**).

We then sought to identify TC and CT synapses formed between the organoids after fusion. To that end, we transduced either the hThO or hCO with hSyn-V5-Mito-APEX2 lentivirus, which localized the V5-tagged peroxidase APEX2 to the mitochondrial matrix in neurons, enabling the identification of the hThO or hCO origin of the presynaptic terminal.^65^ APEX2^+^ hThOs were fused with APEX2^−^ hCOs (or vice versa) to form TC assembloids. Light microscopy and immunolabeling identified V5^+^ puncta that co-localized with neurons expressing hSyn-GFP (**Figure 2I**). Reaction with DAB produced strong contrast in the matrix of APEX2^+^ mitochondria in TEM images. TEM images revealed APEX2^+^ mitochondria in presynaptic terminals from the hThO that formed TC synapses within the hCO after fusion (**Figure 2J**). Conversely, we observed APEX2^+^ mitochondria in presynaptic terminals from the hCOs that formed CT synapses within the hThOs after fusion (**Figure 2K**). This analysis confirmed that assembloids contained both TC and CT synapses.

Whole-cell patch-clamp electrophysiology recordings confirmed that these synapses were functional (**Figure 3**). Electrical synaptic stimulation of the hThO (**Figure 3A**) or hCO (**Figure 3B**) evoked excitatory postsynaptic currents (EPSCs) in cells recorded in the hCO or hThO, respectively. The likelihood of evoking a synaptic response varied among assembloids (**Figure 3A** and **3B**); on average, the chance of cells responding to electrical stimulation of the opposite organoid was 61% for the TC synapses and 58% for the CT synapses (**Figure 3B**). To begin characterizing this synaptic response, we calculated the paired-pulse ratio (PPR) of EPSCs, a classic measure of presynaptic short-term plasticity, at TC and CT synapses.^66–72^ In response to a pair of stimuli applied to a presynaptic neuron, TC synapses elicited paired-pulse depression, wherein the second postsynaptic response was weaker than the first. In contrast, CT synapses were more prone to paired-pulse facilitation, wherein the second response was stronger than the first (**Figure 3C**). Both results resemble previous observations of these synapses in animal models.^66–72^

**Figure 3.**
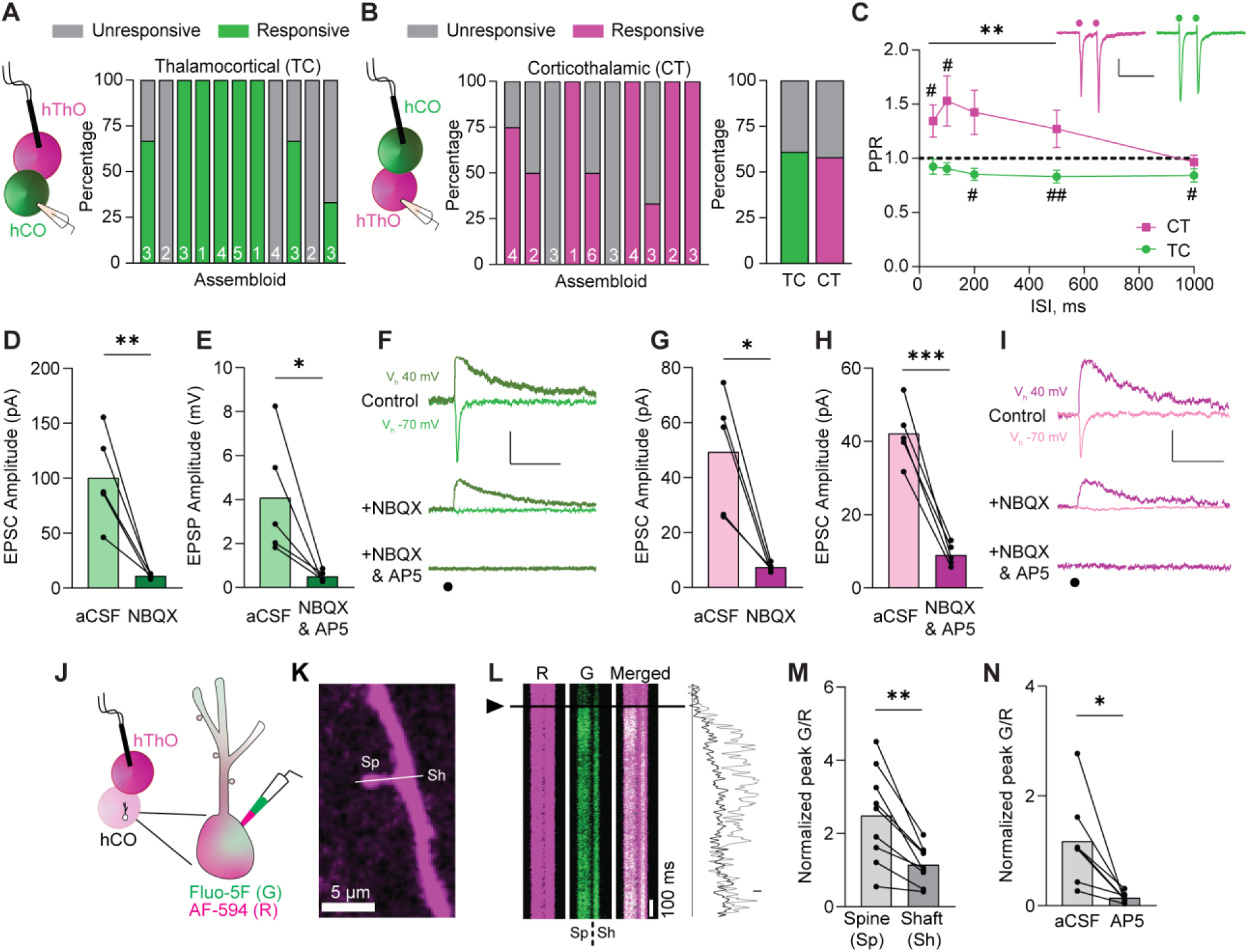
Assembloids contain glutamatergic TC and CT synapses. (**A**) Left: Schematic of the recording configuration for the TC pathway. Right: Bar graph of the percentage of responsive (green) and unresponsive (gray) cells in 11 assembloids. The numbers of cells recorded per assembloid are shown in the bars. (**B**) Left: Schematic of the recording configuration for the CT pathway. Middle: The percentages of hThO cells that responded (magenta) or did not respond (gray) to hCO stimulation across 10 assembloids. Right: Bar graph of the average percentage of responsive cells for TC and CT synapses, based on (**A**) and (**B**). (**C**) Line graph of PPRs across five interstimulus intervals (ISIs) in CT (magenta) and TC (green) synapses [one-sample *t*-test: µ = 1, #p <0.05, ##p <0.01, n = 18-23 cells/9-13 assembloids (TC), n = 8-24/7-12 (CT)]. Differences between CT and TC synapses were evaluated by unpaired *t*-test (**p <0.01). Inset: Sample traces depicting PPRs in CT and TC synapses. Circles represent stimulus artifacts. (**D**) Average TC EPSC amplitude [holding potential (Vh) –70 mV] in the presence of NBQX (3 µM) is significantly decreased compared to control aCSF conditions (paired *t*-test: **p = 0.009, n = 5 cells/2 assembloids). (**E**) The average TC EPSP amplitude (Vh +40 mV) in the presence of NBQX and AP5 (50 µM) is significantly lower than in control aCSF (paired *t*-test: *p = 0.038, n = 5 cells/3 assembloids). (**F**) Traces of evoked TC AMPAR- and NMDAR-mediated currents in control aCSF and in the presence of NBQX or NBQX and AP5, respectively. Circle represents the stimulus artifact. (**G**) Average CT EPSC amplitude (Vh –70 mV) in the presence of NBQX is significantly decreased compared to control aCSF conditions (paired *t*-test: *p = 0.012, n = 5 cells/3 assembloids). (**H**) The average CT EPSC amplitudes (Vh +40 mV) are significantly reduced in the presence of NBQX and AP5 compared to control aCSF (paired *t*-test: ***p = 0.0006, n = 5 cells/3 assembloids). (**I**) Example traces of evoked CT AMPAR- and NMDAR-mediated currents in aCSF and in the presence of NBQX or NBQX and AP5, respectively. (**J**) Schematics of two-photon calcium imaging in postsynaptic dendritic spines of hCO cells upon hThO stimulation. Alexa Fluor 594: AF-594 (R), magenta; Fluo-5F (G), green. (**K**) Image of a dendrite of an hCO cell. Line scans (white line) were performed across a dendritic spine (Sp) and parent dendritic shaft (Sh). (**L**) Left: Representative changes in G/R of Sp and Sh responses over time to a single synaptic stimulation (arrowhead and black line). Right: Representative line scans of Sp (light gray) and Sh (dark gray). (**M**) Average changes in synaptically evoked G/R (paired *t*-test: **p = 0.002, n = 9 cells/4 assembloids). (**N**) Average changes in synaptically evoked Sp G/R in aCSF and in the presence of AP5 (paired *t*-test: *p = 0.018, n = 7 cells/5 assembloids). Data in (**D**), (**E**), (**G**), (**H**), (**M**), and (**N**) are shown as the mean values with individual responses overlaid. Grouped data (**C**) are shown as mean ± SEM. Scale bars (**C**): 20 pA, 200 ms. Scale bars (**F**), (**I**): 40 pA, 100 ms. Scale bar (**L**): 20% G/R. See **Figure S7** for snRNA-seq data supporting glutamatergic communication between hThO and hCOs.

Next, we used whole-cell patch-clamp electrophysiology to further characterize TC and CT synaptic transmission. Evoked TC and CT EPSCs showed typical glutamatergic ionotropic properties comprising a fast α-amino-3-hydroxy-5-methyl-4-isoxazoleproprionic acid receptor (AMPAR)-mediated component blocked by the AMPAR inhibitor NBQX (3 µM) [86.7% ± 3.3% AMPAR current reduction for TC synapses (**Figure 3D** and **3F**); 82.2% ± 3.8% AMPAR current reduction for CT synapses, (**Figure 3G** and **3I**)] and a slow *N*-methyl-D-aspartate receptor (NMDAR)-mediated component blocked by the NMDAR inhibitor AP5 (50 µM) [84.0% ± 4.6% NMDAR current reduction for TC synapses (**Figure 3E** and **3F**); 78.4% ± 2.9% NMDA current reduction for CT synapses (**Figure 3H** and **3I**)].

Using whole-cell patch-clamp electrophysiology and two-photon calcium imaging, we identified the sites of synaptic transmission within a dendritic tree. We detected synaptically evoked calcium transients in hCO cells upon stimulation of hThOs (**Figure 3J-3M**). These postsynaptic sites in the hCO cells resembled dendritic spines described in cortical neurons of animal models (**Figure 3K**).^73^ Stimulation of the hThO evoked stronger calcium transients in dendritic spines compared to that in parent dendritic shafts (**Figure 3L** and **3M**), which suggested that the dendritic spines were synaptically connected to hThO axons. Moreover, calcium transients in dendritic spines were blocked by AP5 [82.4% ± 5.2% reduction (**Figure 3N**)], indicative of glutamatergic synaptic transmission.

### TC and CT synapses undergo LTP in assembloids

Having established the existence of functional synaptic connections between hThOs and hCOs, we tested whether the TC and CT pathways undergo long-term synaptic plasticities, specifically LTP and LTD. To examine whether TC synapses undergo LTP, we tested several LTP-induction protocols: short and long spike-timing–dependent plasticity (STDP) induction protocols and high-frequency tetanization. TC LTP was reliably induced by high-frequency (40-Hz) tetanization of thalamic inputs (**Figure 4A**), which increased EPSC amplitudes by 168.2% ± 19.3% compared to baseline (**Figure 4B** and **4C**). STDP is induced by stimulating presynaptic inputs and directly depolarizing the postsynaptic cell **(Figure 4D** and **4G**); STDP is based on the precise order and timing of pre- and postsynaptic activity.^74–76^ Following a short (×1) STDP-induction protocol, the amplitude of TC EPSCs increased by 144.4% ± 17.8%, compared to baseline (**Figure 4D-4F**). Following a long (×3) STDP-induction protocol, EPSC amplitudes increased by 223.9% ± 24.4%, compared to baseline (**Figure 4G-4L**). This TC LTP was observed in all tested (9/9) cells, from six separate assembloids, across three batches of differentiation (**Figure 4H**). TC LTP was not caused by changes in series resistance (**Figure 4I**); thus, it represents a true activity-dependent potentiation of synaptic strength.

**Figure 4.**
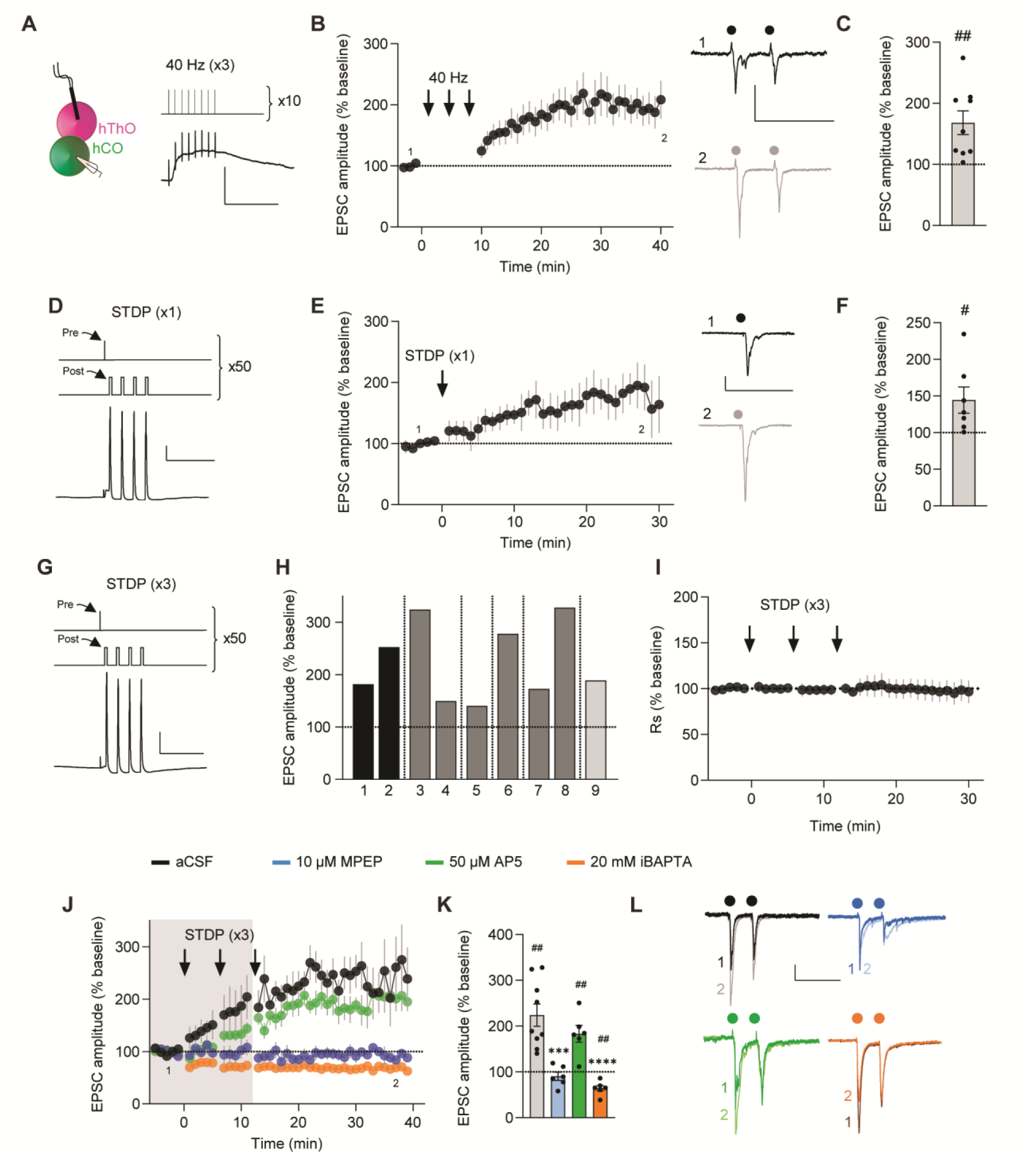
TC synapses in assembloids undergo LTP via multiple protocols. (**A**) Left: Schematic of the recording configuration. Right, top: 40-Hz electrical stimulation LTP-induction protocol. Right, bottom: Representative trace of a response to 10-Hz stimulation. (**B**) Left: Time course data demonstrating that 40-Hz stimulation repeated three times (arrows) induces LTP in TC assembloids (n = 9 cells/9 assembloids). Right: Representative traces from the first 5 min (1, dark) and final 5 min (2, light) of the experiment. Circles indicate electrical stimuli. (**C**) Bar graph of group data after 40-Hz induction from (**B**) shows EPSC amplitudes significantly differ from baseline values (one-sample *t*-test, µ = 100, ##p = 0.0077). (**D**) Top: Spike-timing–dependent plasticity (STDP) was induced by stimulating presynaptic hThO inputs (Pre) and then delivering four current injections (2-nA) to the postsynaptic cell (Post), repeated 50 times. Bottom: Representative trace of an hCO cell’s response to stimulation and depolarization. (**E**) Left: Time course data demonstrating that the short (×1) STDP protocol (arrow) in TC assembloids induces LTP (n = 7 cells /3 assembloids). Right: Representative traces from the first (dark) and final (light) 5 min of the experiment. (**F**) Bar graph of group data following the ×1 STDP induction from (**E**) shows that EPSC amplitudes significantly differ from baseline values (one-sample *t*-test, µ = 100, #p = 0.04). (**G**) Top: Long (×3) STDP-induction protocol, as in (**D**) but repeated three times every 5 min. Bottom: Representative trace of a response to stimulation and depolarization. (**H**) Bar graph showing the average responses from nine cells from six assembloids after TC LTP induction. Shades of gray indicate different batches of assembloids; vertical lines denote separate assembloids. (**I**) Time course of series resistance (Rs) normalized to the 5-min baseline period demonstrating TC LTP is not due to changes in Rs. (**J**) Time course demonstrating the ×3 STDP protocol (arrows) induces LTP in TC synapses (black, n = 9 cells/6 assembloids). MPEP (blue, n = 6 cells/5 assembloids) or iBAPTA blocked LTP (orange, n = 6 cells/4 assembloids). AP5 did not block TC LTP (green, n = 6 cells/3 assembloids). Shaded area depicts the presence of bath-applied drugs. (**K**) Bar graph of group data following ×3 STDP induction from (**J**). Differences from baseline were evaluated by one-sample *t*-test (µ = 100, ##p <0.01). Differences between treatments and aCSF were evaluated by one-way ANOVA, p <0.0001. Dunnett’s test: ***p =0.0001, ****p <0.0001. (**L**) Example traces from the first (1) and final (2) 5 min of the experiment across conditions. Scale bars for (**A**): 20 mV, 200 ms. Scale bars for (**B**), (**E**): 50 pA, 200 ms. Scale bars for (**D**), (**G**): 40 mV, 100 ms. Scale bars for (**L**): 20 pA, 200 ms. Data shown are mean ± SEM (**B**), (**E**), (**I**), and (**J**), with individual data points overlaid as dots in (**C**), (**F**), and (**K**). For (**B**), (**E**), (**J**), and (**L**) the first (1) and final (2) 5 min of the experiment are noted. See **Figure S7** for analysis of paired-pulse ratio (PPR) measures and analysis of organoid/assembloid age and TC LTP expression.

Next, we investigated the mechanisms underlying TC LTP in assembloids. Bath application of the metabotropic glutamate 5 (mGluR5) antagonist MPEP (10 µM) blocked TC LTP, but the NMDAR antagonist AP5 (50 µM) did not (**Figure 4J** and **4K**). TC LTP also required postsynaptic Ca^2+^. When we included the Ca^2+^ chelator BAPTA (20 mM) in the internal pipette solution (iBAPTA), the long STDP protocol not only failed to induce LTP but also moderately reduced the TC EPSC amplitude (**Figure 4J** and **4K**). We then tested if TC LTP was expressed presynaptically by measuring changes in PPR after LTP induction. PPR decreased (suggesting an increase in the probability of glutamate release from presynaptic terminals) compared to baseline in control and AP5 conditions, and this change was blocked in the presence of MPEP (**Figure S7A** and **S7B**). Together, these findings demonstrate that the TC pathway in assembloids undergoes LTP via multiple induction protocols, with the long (×3) STDP-evoked LTP induced postsynaptically and expressed, at least partially, presynaptically through mGluR5-dependent mechanisms.

CT synapses also underwent LTP after the long STDP-induction protocol (**Figure 5**). LTP was observed in 12/14 cells recorded from nine separate assembloids across two differentiation batches (**Figure 5B**). On average, EPSC amplitudes increased by 158.3% ± 15.2%, compared to baseline after LTP induction (**Figure 5C** and **5E**). CT LTP was not caused by changes in series resistance (**Figure 5D**), representing a true potentiation of synaptic strength. Inclusion of iBAPTA in the internal pipette solution blocked CT LTP (**Figure 5C** and **5E**), and unlike TC LTP, CT LTP was also blocked by separate application of MPEP and AP5 (**Figure 5C** and **5E**). There were no changes in PPR in any of the conditions (**Figure S7C** and **S7D**), suggesting that CT LTP does not involve changes in presynaptic-release probabilities. These data suggest that the long (×3) STDP induction protocol generates LTP in the CT pathway in assembloids, and this LTP is induced and expressed postsynaptically through both mGluR5- and NMDAR-dependent mechanisms.

**Figure 5.**
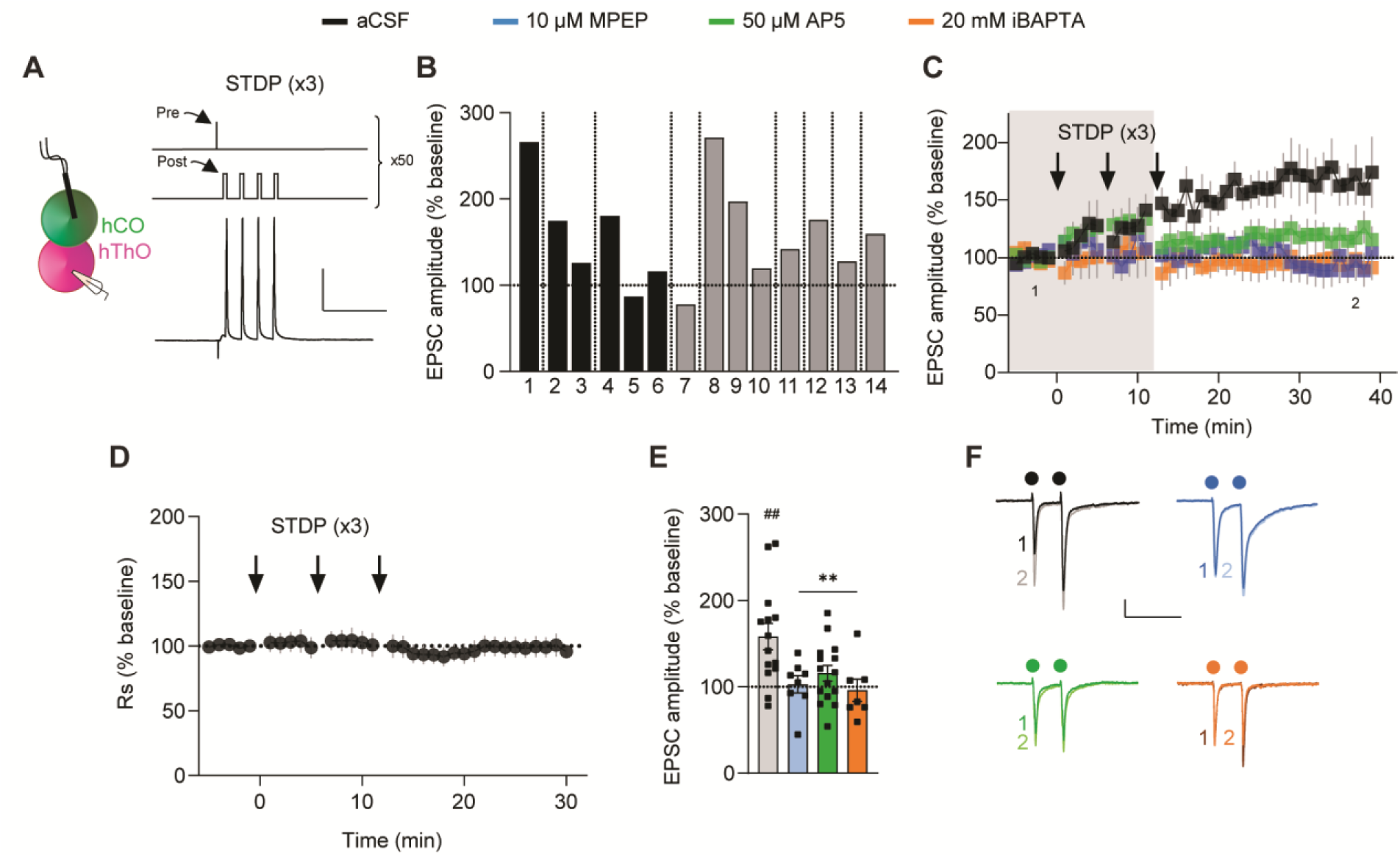
CT synapses in assembloids undergo LTP. (**A**) Left: Schematic of the recording configuration to induce CT LTP. Right: The long (×3) STDP-induction protocol and example response. (**B**) Bar graph showing the average responses in 14 cells from 9 assembloids after CT LTP induction in aCSF. Shades of gray indicate different batches of assembloids; vertical lines denote separate assembloids. (**C**) Time course demonstrating that ×3 STDP delivery (arrows) induces LTP in CT synapses (black, n = 14 cells/9 assembloids). MPEP (blue, n = 8 cells/6 assembloids), AP5 (green, n = 15 cells/7 assembloids), or iBAPTA (orange, n = 7 cells/3 assembloids) blocked LTP. Shaded area depicts the presence of bath-applied drugs. The first (1) and final (2) 5 min of the experiment are noted. (**D**) Time course of Rs demonstrating that CT LTP is not due to changes in Rs. (**E**) Bar graph of group data from (**C**). Differences from baseline were evaluated by one-sample t-test (µ = 100, ##p <0.01). Differences between treatments and aCSF were evaluated by one-way ANOVA, p = 0.0053. Dunnett’s test: **p <0.01. (**F**) Example traces from the first (1) and final (2) 5 min of the experiment across conditions. Circles indicate electrical stimulation. Scale bars for (**A**): 40 mV, 100 ms. Scale bars for (**F**): 50 pA, 200 ms. Data shown are mean ± SEM (**C**), (**D**) with individual data points overlaid in (**E**). See **Figure S7** for PPR analysis and analysis of organoid/assembloid age and CT LTP expression.

### TC and CT synapses undergo LTD in assembloids

The activity-dependent weakening of synaptic transmission between brain regions through LTD is a key component of learning.^77^ Therefore, we tested whether TC and CT synapses in assembloids undergo LTD. Delivering low-frequency (1-Hz) stimulation to the hThO while recording from hCO cells depressed EPSCs in 8/9 cells recorded in nine assembloids from three differentiation batches (**Figure 6A** and **6B**). On average, TC EPSC amplitudes decreased by 59.4% ± 8.1% of baseline after low-frequency stimulation (**Figure 6C-6F**). TC LTD was blocked by iBAPTA or bath-applied MPEP or AP5 (**Figure 6C** and **6E**). Low-frequency stimulation of the hThO did not change PPR across any of the conditions (**Figure S7E** and **S7F**). These findings provide evidence for a postsynaptically induced and expressed LTD in the TC pathway that depends on both mGluR5s and NMDARs.

**Figure 6.**
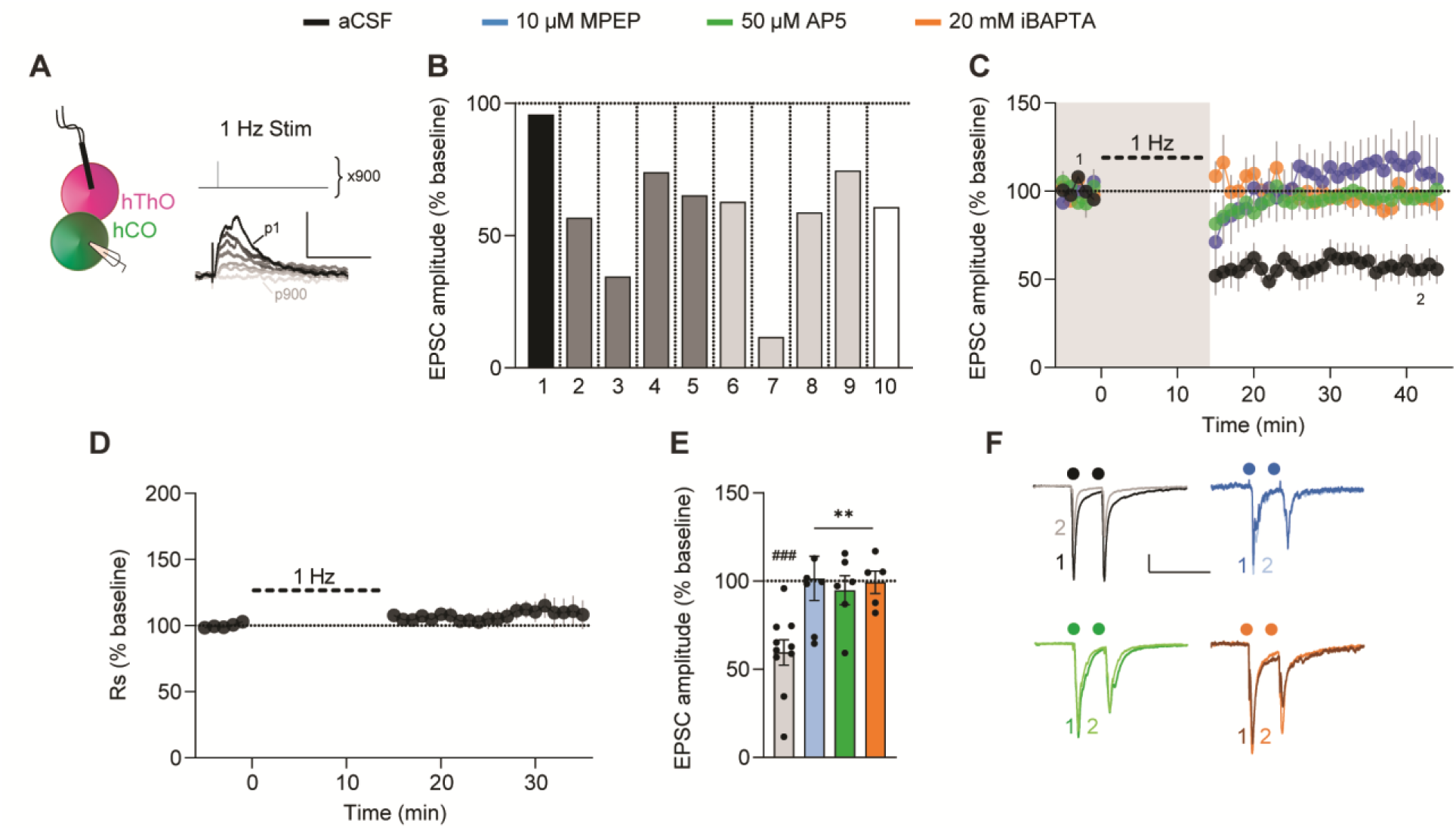
TC synapses in assembloids undergo LTD. (**A**) Left: Schematic of the recording configuration to induce TC LTD. Right, top: LTD was induced with electrical stimulation delivered at 1 Hz for 900 pulses. Bottom: Example responses of a cell to a subset of the 900 pulses, dark-to-light traces depict the responses as the number of pulses progressed (from pulse (p) 1 to p900). (**B**) Bar graph showing the average responses from 10 individual cells from 10 assembloids after TC LTD induction in aCSF. Shades of gray indicate different batches of assembloids; vertical lines denote separate assembloids. (**C**) Time course data demonstrating that 1-Hz electrical stimulation (thick dashed line) induces LTD in TC synapses (black, n = 10 cells/10 assembloids). MPEP (blue, n = 7 cells/5 assembloids), AP5 (green, n = 6 cells/5 assembloids), or iBAPTA (orange, n = 5 cells/4 assembloids) blocked LTD. Shaded area depicts the presence of bath-applied drugs. The first (1) and final (2) 5 min of the experiment are noted. (**D**) Time course of Rs normalized to the 5-min baseline period, demonstrating that TC LTD is not due to changes in Rs. (**E**) Bar graph of group data after 1-Hz stimulation from (**C**). Differences from baseline were evaluated by one-sample *t*-test (µ = 100, ###p <0.005). Differences between treatments and aCSF were evaluated by one-way ANOVA, p = 0.0046. Dunnett’s test: **p <0.01. (**F**) Example traces from the first (1) and final (2) 5 min of the experiment across conditions. Circles indicate electrical stimulation. Scale bars for (**A**): 5 mV, 100 ms. Scale bars for (**F**): 40 pA, 200 ms. Data shown are mean ± SEM (**C**), (**D**), and (**E**), with individual data points overlaid in (**E**). See **Figure S7** for PPR analysis and analysis of organoid/assembloid age and TC LTD expression.

Low-frequency stimulation of hCO inputs to hThO cells also reliably induced LTD, as it was observed in 10/10 cells recorded in nine assembloids across two batches of differentiation (**Figure 7A** and **7B**). On average, the expression of CT LTD was reflected in a 65.8% ± 5.2% reduction of baseline EPSC amplitudes (**Figure 7C-7F**). CT LTD was blocked by iBAPTA or bath application of MPEP or AP5 (**Figure 7C** and **7E**). PPR at CT synapses was unchanged after 1-Hz stimulation across all conditions (**Figure S7G** and **S7H**). Neither TC LTD nor CT LTD occurred due to changes in series resistance (**Figures 6D** and **7D**, respectively). These data suggest that CT LTD is induced and expressed postsynaptically and requires both mGluR5s and NMDARs Notably, reversing the order of presynaptic stimulation and postsynaptic depolarization, in a reverse long STDP protocol, did not induce LTD in either the TC or CT pathways (data not shown). The mechanisms underlying TC and CT LTP/LTD were supported by snRNA-seq experiments, which revealed that neurons in hThOs and hCOs contain transcripts encoding group 1 mGluRs (predominantly *GRM5*, which encodes mGluR5), NMDAR subunits, and AMPAR subunits, including the direct targets of MPEP and AP5 (**Figure S7I** and **S7J**). Finally, neither the age of the individual organoid nor days post-fusion of the assembloid affected the expression of TC or CT LTP/LTD (**Figure S7K**). The number of batches of individual organoids used for each experimental condition is listed in **Table S2**.

**Figure 7.**
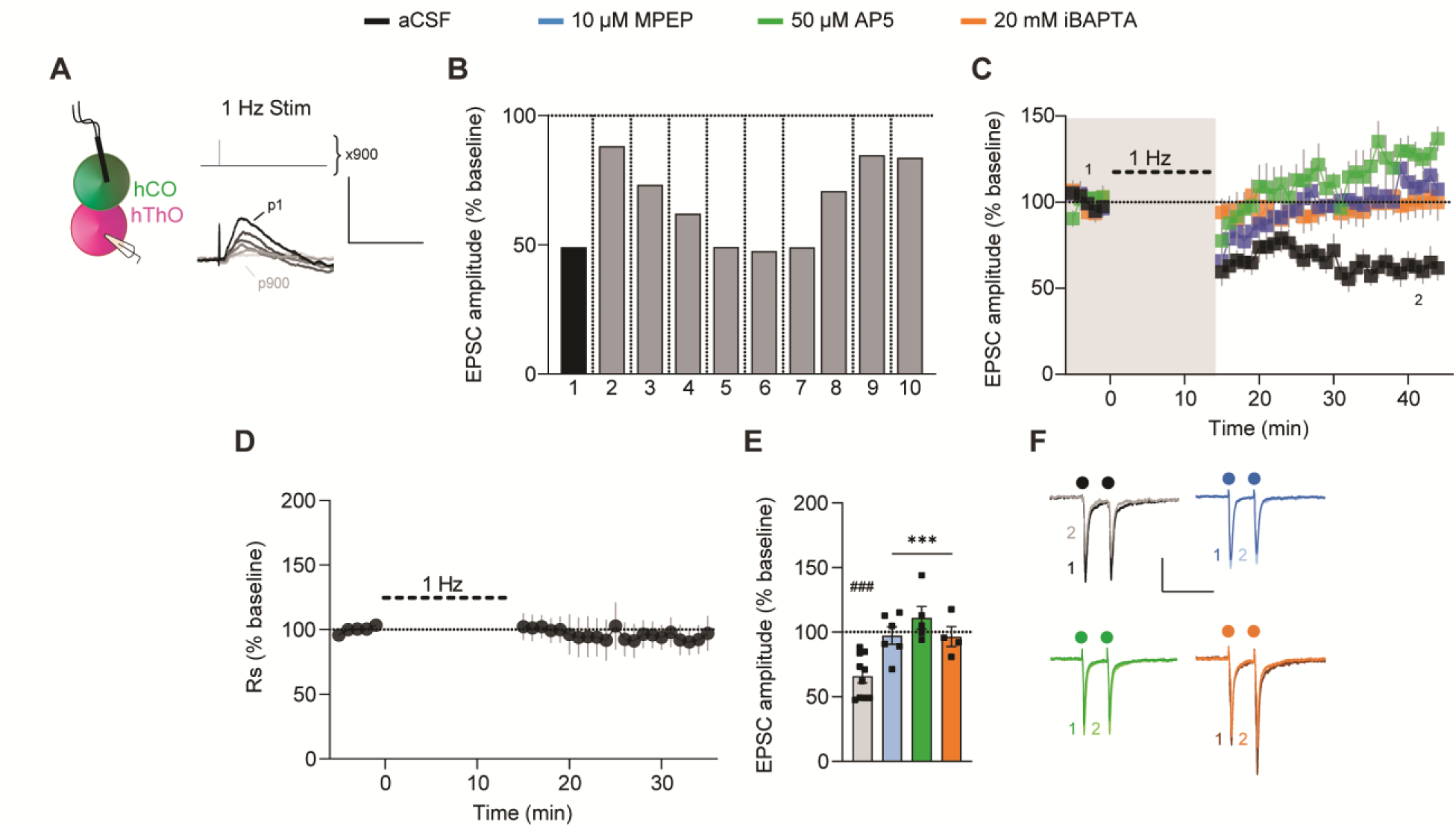
CT synapses in assembloids undergo LTD. (**A**) Schematic of the experimental condition for CT LTD induction and the 1-Hz LTD-induction protocol. (**B**) Bar graph of the average responses of 10 individual cells from 9 assembloids after CT LTD induction in aCSF. Shades of gray indicate different batches of assembloids; vertical lines denote separate assembloids. (**C**) Time course data show that 1-Hz stimulation (thick dashed line) induced LTD in CT synapses (black, n = 10 cells/9 assembloids). MPEP (blue, n = 6 cells/6 assembloids), AP5 (green, n = 5 cells/4 assembloids), or iBAPTA (orange, n = 4 cells/4 assembloids) blocked LTD induction. Shaded area depicts the presence of bath-applied drugs. The first (1) and final (2) 5 min of the experiment are noted. (**D**) Time course of Rs normalized to the 5-min baseline period, demonstrating that CT LTD is not due to changes in Rs. (**E**) Bar graph of group data after 1-Hz stimulation from (**C**). Differences from baseline were evaluated by one-sample *t*-test (µ = 100, ###p <0.001). Differences between treatments and aCSF were evaluated by one-way ANOVA, p = 0.0004. Dunnett’s test: ***p <0.001. (**F**) Example traces from the first (1) and final (2) 5 min of the experiment across conditions. Circles indicate electrical stimulation. Scale bars for (**A**): 5 mV, 100 ms. Scale bars for (**F**): 50 pA, 200 ms. Data shown are mean ± SEM (**C**), (**D**), and (**E**), with individual data points overlaid in (**E**). See **Figure S7** for PPR analysis and analysis of organoid/assembloid age and CT LTD expression.

## DISCUSSION

Here we describe a novel, hiPSC-derived TC assembloid system for exploring synaptic transmission and synaptic plasticity in human neural circuits. Within assembloids, hThOs and hCOs form reciprocal glutamatergic synapses capable of short- and long-term synaptic plasticity. Previous work describing synaptic plasticity in organoids has exclusively relied on multielectrode array (MEA) recordings,^78^ which measure extracellular spike and local field activities that arise from a large, heterogeneous group of cells. Reflecting this heterogeneity, MEA recordings in organoids detected LTP, LTD, and bidirectional short-term synaptic plasticity in response to identical induction protocols.^78^ Our study used whole-cell patch-clamp electrophysiology, a well-validated method that has enabled decades of synaptic physiology discoveries in multiple species. By accurately measuring electrically evoked synaptic currents in single postsynaptic cells, we found that the vast majority (92.9%) of synaptically connected cells underwent LTP or LTD during the respective induction protocols. Furthermore, these synapses displayed a degree of specificity, as one established induction protocol (reverse long STDP protocol) failed to induce LTD at either synapse. Together, our findings suggest that synaptic plasticity is robust, highly replicable, and selective in the TC assembloid system.

Few studies have examined synaptic transmission within a specific circuit in human brain organoids.^29–31^ More commonly, the properties of spontaneously released neurotransmitters onto single cells in organoids have been characterized,^79–81^ but the presynaptic sources of that transmission were not defined. Based on our snRNA-seq data, NeuronChat analysis, and whole-cell patch-clamp electrophysiological recordings, we believe the TC recordings arise from glutamatergic cortical neurons receiving synaptic inputs from glutamatergic thalamic neurons, while CT recordings arise from glutamatergic thalamic neurons receiving synaptic inputs from glutamatergic cortical neurons. However, one key limitation of this study is the uncertainty of which specific ExN clusters comprise the pre- and postsynaptic cell populations. In defined, heterogeneous systems such as the rodent cortex, multiple modes of identification are used to define neuronal subtypes, e.g. laminar location, cellular morphology, electrophysiological profiles, and genetics. However, in the heterogeneous human-derived organoid system, most of these characteristics remain poorly defined. Patch-seq, which couples single-cell electrophysiological recordings with single-cell RNA-seq and morphology data,^82^ would directly address the question of intra-organoid cell type specificity. However, Patch-seq has not yet been established in organoids, in studies of long-term synaptic plasticity, or in fetal brain tissue. Therefore, we believe this is a very exciting and powerful future direction for the organoid and developmental neurobiology fields.

Although there are some similarities between our LTP/LTD findings from human assembloids and those from rodents, specifically the high prevalence for NMDAR-mediated LTP and LTD,^83^ in general the mechanisms underlying LTP/LTD in the rodent TC and CT pathways are distinct from what we report here.^84–86^ In mouse auditory and somatosensory TC pathways, LTP depends on postsynaptically expressed group 1 mGluRs, whereas LTP in the barrel cortex requires postsynaptic NMDAR activation and subsequent Ca^2+^ entry into the postsynaptic cell.^87–91^ CT LTP in rodents is expressed presynaptically, requires a rise in presynaptic Ca^2+^ and protein kinase A activation,^92^ but does not require the activation of NMDARs or mGluRs.^92^ In mice, LTD within the somatosensory cortex is mediated by NMDARs,^93^ and LTD in the barrel cortex requires presynaptic type 1 cannabinoid receptors.^89^ In contrast, we found that three of the four types of long-term synaptic plasticity we measured required both mGluR5 and NMDARs. A functional link between group 1 mGluRs and NMDAR activity is seen in various brain regions.^94,95^ In mouse cortical neurons, the activation of mGluR1 potentiates NMDAR-mediated currents through downstream signaling.^96^ Given our findings, a similar mGluR5-dependent mechanism may exist in human-derived TC assembloids.

Our results may reflect species-specific differences between rodents and humans in the expression and maintenance of long-term synaptic plasticity. However, organoids resemble fetal human brain more closely than they do postnatal structures,^97^ and most rodent studies of synaptic physiology are conducted in postnatal animals. Unlike the adult counterparts, the immature thalamus does not contain well-differentiated nuclei, and the fetal cortical plate lacks well-defined layers. Organoids in their current forms display similar limitations in their structural organization. Therefore, the mechanisms we define in TC assembloid synaptic plasticity may differ from those observed in rodents due to species differences, developmental differences, or a combination thereof.

The organoid field is constantly improving. A recent report described hThOs that more closely resemble the thalamic reticular nucleus.^98^ Future studies will derive organoids that better model specific projection nuclei of the thalamus. In the developing brain, thalamic inputs mold the laminar, columnar, and functional organization of the cortex.^99,100^ Similarly, more mature thalamic projections might promote organizational maturation in hCOs, and more organized TC assembloids might better model the diversity of synaptic plasticity mechanisms observed across TC sensory pathways. Assembloids modeling other brain structures^101–103^ (or synaptic targets outside the brain)^104^ might also elucidate mechanisms that differ between neural circuits.

We anticipate that hiPSC-derived organoids and assembloids will provide a particularly useful model system for exploring synaptic pathology in human neurologic and psychiatric disorders. To date, most organoid studies have focused on disease-associated changes in gene expression, cellular composition, or neural network activity. Our data suggest that assembloids derived from patient hiPSCs or from hiPSCs carrying disease-associated mutations can be used to model disease-associated deficits in synaptic transmission and synaptic plasticity. We expect that TC assembloids will be particularly useful in this respect, as functional abnormalities in many thalamic nuclei are linked to psychiatric disorders, including schizophrenia.^105^ The findings we present here provide a foundation for those future studies.

## Acknowledgments

We thank Yiping Fan, Dale Hedges, and Daniel Estevez Prado (St. Jude Center for Applied Bioinformatics, Transcriptomics Group) for assistance with bulk RNA-seq data analysis; Lawrence Reiter for providing the TP-190a and TP-189 dental pulp stem cells for reprogramming; Sergiu Pasca for providing the 2242, 1205, and 8858 hiPSC lines; Angela McArthur for manuscript editing; and Zakharenko lab members for constructive comments.

## Author contributions

Conceptualization: MHP, KTT, SSZ

Electrophysiology experiments: MHP, ITB, YJ

Molecular biology experiments: KTT, AN, ABS

Two-photon imaging experiments: ITB

hiPSC line maintenance, organoid differentiation, and assembloid preparation: AN, KDN

Electron microscopy: NBK, CGR

snRNA-seq sample preparation: AJT, JBB

snRNA-seq data analysis: CAR, KTT

Design and production of hiPSC reporter lines: STP, SMP-M

Visualization: MHP, KTT

Funding acquisition: SSZ

Supervision: SSZ

Writing – original draft: MHP, KTT

Writing – review & editing: SSZ

## Data and materials availability

The snRNA-seq data that support the findings of this study have been deposited in SRA under the Bioproject ID PRJNA999219. Bulk RNA-seq data have been deposited in SRA under Bioproject ID PRJNA1001283. Code used for the analysis of snRNA-seq data is available at Github (https://github.com/ZakharenkoLab/Thalamic_and_Cortical_Organoid_snRNASeq). Additional R code is available upon request.

## Supplemental Materials

Figures S1 to S7

Table S1 and S2

## STAR Methods

### Human iPSC culture

The use of hiPSCs for the generation of organoids was approved by the St. Jude Institutional Review Board. TP-190a and TP-189 were derived from dental pulp cells from neurotypical male and female subjects, respectively, with normal karyotypes. Cells were reprogrammed using episomal plasmids (ALSTEM LLC). The 2242 (i.e., 2242-1), 1205 (i.e., 1205-4), and 8858 (i.e., 8858-3) lines/clones were previously published.^106^

All hiPSC lines were maintained in culture on human ES-qualified Matrigel (5264004, Corning) in complete mTeSR Plus (100-0276, STEMCELL Technologies) at 5% O_2_, 37 °C, and 5% CO_2_. The cultures were passaged with Versene (15040066, ThermoFisher). Genetically modified reporter hiPSC lines were validated before differentiation (**Figure S1**). Specifically, six assays were performed: (1) Colonies were immunostained for six pluripotency markers^107^ (Stemlight Pluripotency Antibody Kit 9656S, Cell Signaling); (2) Expression of five additional pluripotency markers^107^ was assayed using a Custom TaqMan RT-qPCR assay designed in-house (manufactured by ThermoFisher); (3) G-banding; (4) Copy number variation at the seven most frequently aberrant chromosomal locations^108–110^ was assayed using a custom TaqMan RT-qPCR assay designed in-house (manufactured by ThermoFisher); (5) Global methylation analysis (Infinium MethylationEPICv1.0 850K Beadchip) was performed to identify methylation status at select epigenetic markers,^111,112^ and results were then compared to established and previously published hiPSC lines;^113^ (6) Trilineage assay (STEMdiff Trilineage Differentiation Kit 05230, STEMCELL Technologies) was performed, and markers of interest^107^ were analyzed using a Custom TaqMan RT-qPCR assay designed in-house (manufactured by ThermoFisher).

### Generating reporter lines

Genome-edited TP-190a hiPSCs were generated using CRISPR-Cas9 technology. Briefly, hiPSCs were pretreated with StemFlex (Thermo-Fisher Scientific) supplemented with 1× RevitaCell (ThermoFisher Scientific) for 1 h. Then, approximately 10^6^ cells were transiently transfected with precomplexed ribonuclear proteins consisting of 250 pmol chemically modified single-guide RNA (sgRNA; Synthego), 165 pmol Cas9 protein (St. Jude Protein Production Core), 500 ng pMaxGFP (Lonza), and 3 µg ssODN donor (for deletion) or 1 µg dsDNA donor (for tagging) via nucleofection (Lonza, 4D-Nucleofector™ X-unit) using solution P3 and program CA-137 in a large (100-µL) cuvette, according to the manufacturer’s recommended protocol. Five days postnucleofection, cells were single-cell sorted by FACS for GFP^+^ (transfected) cells and plated onto Vitronectin XF (Stem Cell Technologies)-coated plates into prewarmed (37°C) StemFlex media supplemented with 1× CloneR (Stem Cell Technologies).

Clones were screened for the desired modification via targeted deep sequencing using gene-specific primers with partial Illumina adapter overhangs on a Miseq Illumina sequencer, as previously described,^114^ or by junction PCR followed by sequencing. Briefly, cell pellets of approximately 10,000 cells were lysed and used to generate gene-specific amplicons with partial Illumina adapters in PCR#1. Amplicons were indexed in PCR#2 and pooled with targeted amplicons from other loci to create sequence diversity. Additionally, 10% PhiX Sequencing Control V3 (Illumina) was added to the pooled amplicon library prior to running the sample on a Miseq Sequencer System (Illumina) to generate paired 2 × 250-bp reads. Samples were de-multiplexed using the index sequences, fastq files were generated, and next-generation sequencing (NGS) analysis of clones was performed using CRIS.py.^115^ Correctly edited clones were identified, expanded, and sequence confirmed. Final clones were authenticated using the PowerPlex® Fusion System (Promega), which was performed at the St. Jude Hartwell Center and tested for mycoplasma by using the MycoAlertTMPlus Mycoplasma Detection Kit (Lonza). Editing-construct sequences and relevant primers are listed in **Table S1**.

### Thalamic organoid generation

The hThOs were generated from TCF7L2-tdTomato reporter hiPSC lines, except for indicated organoids in **Figure S2**. For differentiation, cryovials were plated and maintained in culture in mTeSR1 (85850, STEMCELL Technologies). At 80% confluency, they were dissociated into single cells with Accutase (AT-104, Innovative Cell Technologies) and plated into low-attachment 96-well V-bottom plates (MS-9096VZ, Sbio) at 10,000 cells/well, in gfCDM media (1:1 IMDM (12440053, Thermofisher): Ham’s F12 (12-615F, Lonza), 1× lipid concentrate (11905031, Thermofisher), 1× antibiotic-antimycotic (15240062, Gibco), 450 µM monothioglycerol (M6145, Sigma), 15 µG/mL apotransferrin (T1428, Sigma), 5 mg/mL BSA (50-255-465, Fisher Scientific) supplemented with 5 µM SB-431542 (TGFβ inhibitor, 1614, Tocris), 1 µg/mL insulin (I9278, Sigma), 1% v/v growth factor–reduced (GFR) Matrigel (354230, Corning), 2 µM thiazovivin (72254, STEMCELL Technologies). On Day (D) 2, half the media was replaced with the same media supplemented with 4 µM dorsomorphin (3093, Tocris). On D4 and D6, half of the media was replaced with gfCDM supplemented with 5 µM SB-431542, 100 nM Smoothened agonist (73414, STEMCELL Technologies), and 20 ng/mL Fgf8b (100-25, PeproTech). In some differentiations, Matrigel was added on D2 instead of D0, but no difference was detected in the resulting organoids. On D8, D10, and D12, three-fourths of the media was replaced with the same but further supplemented with 30 ng/mL BMP7 (354BP010, R&D Systems) and 1 µM MEKi PD0325901 (S1036, R&D Systems). On D14, D16, and D18 half the media was replaced with the same, but gfCDM was substituted with thalamic N2 media (DMEM:F12, 10% ES-FBS (ES-009-C, SIGMA), 1× N2 supplement (17502-048, Gibco), 1× Glutamax (35050061, Gibco), and 1× antibiotic-antimycotic.

On D20, all organoids were transferred to a magnetic stir bioreactor (BWS-S03N0S-6, ABLE Corporation, Tokyo) in thalamic N2 media and agitated at 40 rpm. On D22 and D24, half the media was replaced with thalamic N2 supplemented with 1× B27 without vitamin A (12587-010, Gibco), 20 ng/mL heat-stable bFGF (PHG0367, ThermoFisher) and 20 ng/mL EGF (AF-100-15-100UG, Peprotech). On D26 and D28, half the media was replaced with the same, but the concentrations of bFGF and EGF were reduced to 10 ng/mL. On D30 and D32, all the media was replaced with thalamic N2 supplemented with 1× B27 without vitamin A.

Starting D35, full media was replaced every 4 days with BrainPhys (05790, STEMCELL Technologies) supplemented with 1× N2, 1× B27 without vitamin A, 10% ES-FBS, 10 ng/mL BDNF (450-02, Peprotech), and 10 ng/mL GDNF (450-10, Peprotech). Starting D70, all the media was changed to BrainPhys supplemented with 1× N2, 1× B27 without vitamin A, 1× glutamax, 1× NEAA (11140050, Gibco), 1× antibiotic-antimycotic, 200 µM ascorbic acid (A4403, Sigma), 100 µM dibutyryl cAMP (D0627, Sigma), 1% ES-FBS, 10 µM DAPT (2634, Tocris), 20 ng/mL BDNF, and 20 ng/mL GDNF. Starting D82, the concentration of BDNF and GDNF was reduced to 10 ng/mL. In addition, after D30, large organoids were pinched into two halves by using a pair of ultra-fine clipper scissors (15300-00, Fine Science Tools) every 5–7 days to avoid large necrotic centers.

### Cortical organoid generation

The hCOs were generated from VGLUT1-tdTomato reporter iPSC lines. At 80% confluency, cell cultures were dissociated into single cells with Accutase (AT-104, Innovative Cell Technologies), and plated into low-attachment 96-well V-bottom plates (MS-9096VZ, Sbio) at 9000 cells/well, in EB media (DMEM:F12, 20% knockout serum replacement (KSR) (10828, Life Technologies), 3% ES-FBS, 1× Glutamax, 1× β-mercaptoethanol (2020-07-30, Gibco), 1× antibiotic-antimycotic) supplemented with 5 µM SB-431542 (TGFβ inhibitor, 1614, Tocris), 2 µM dorsomorphin (3093, Tocris), 3 µM IWR1e (Wnt inhibitor, 681669, EMD Millipore), 1% v/v GFR-Matrigel, and 2 µM thiazovivin.

In some differentiations, 0% or 0.5% v/v GFR-Matrigel was added on D0 but no difference in the resulting organoids was detected. On D2, half the media was replaced with the same but without Matrigel. On D4 and D6, half the media was replaced with GMEM KSR media (GMEM, 20% KSR, 1× NEAA (Gibco), 1× sodium pyruvate (11360070, Gibco), 1× β-mercaptoethanol, 1× antibiotic-antimycotic) supplemented with 5 µM SB-431542, 3 µM IWR1e, 2.5 µM cyclopamine (72074, STEMCELL Technologies), and 2 µM thiazovivin. On D8, half the media was replaced with GMEM KSR media supplemented with 5 µM SB-431542, 3 µM IWR1e, and 2.5 µM cyclopamine. On D10, D12, D14, and D16, half the media was replaced with GMEM KSR media supplemented with 5 µM SB-431542 and 3 µM IWR1e. On D18 and D20, half the media was replaced with CBO N2 media (DMEM:F12, 1× chemically defined lipid concentrate (11905-031, Life Technologies), 1× N2 supplement (17502-048, Gibco) and 1× antibiotic-antimycotic) supplemented with 1× B27 supplement without vitamin A (12587-010, Gibco), 20 ng/mL heat-stable bFGF, and 20 ng/mL EGF. On D22, organoids were transferred to a magnetic stir bioreactor (BWS-S03N0S-6, ABLE Corporation) in CBO N2 media supplemented with 1× B27 supplement without vitamin A, 20 ng/mL heat-stable bFGF and 20 ng/mL EGF, and agitated at 40 rpm. Half of the media was replaced with the same on D24, D26, and D28. On D30, the media was changed to CBO FBS media (DMEM:F12, 1× chemically defined lipid concentrate (11905-031, Life Technologies), 1× N2 supplement, 10% ES-FBS, 5 µg/mL heparin and 1× antibiotic-antimycotic) supplemented with 1× B27 supplement without vitamin A. Full media was replaced every 4 days. On D42 and D46, the media was changed to CBO FBS media supplemented with 1× B27 supplement without vitamin A, 10 ng/mL BDNF (450-02, Peprotech), and 10 ng/mL GDNF (450-10, Peprotech). Starting D50, all the media was replaced every 4 days with BrainPhys (05790, STEMCELL Technologies) supplemented with 1× N2 supplement, 50× B27 supplement without vitamin A, 10% ES-FBS, 10 ng/mL BDNF and 10 ng/mL GDNF. Starting D70, media was changed to BrainPhys supplemented with 1× N2, 50× B27 without vitamin A, 1× glutamax, 1× NEAA, 1× antibiotic-antimycotic, 200 µM ascorbic acid, 100 µM cAMP, 1% ES-FBS, 10 µM DAPT, 20 ng/mL BDNF, and 20 ng/mL GDNF. Starting at D82, the concentration of BDNF and GDNF was reduced to 10 ng/mL. In addition, after D30, large organoids were pinched into 2 halves by using a pair of ultra-fine clipper scissors every 5–7 days to avoid large necrotic centers.

### Generation of thalamocortical assembloids

Between D90 and D120, TCF7L2-tdTomato^+^ hThOs were paired with VGLUT1-tdTomato^+^ hCOs of similar age. Each pair was transferred to a well of a low-attachment, 24-well plate in 500 mL Fusion media (BrainPhys supplemented with 1× N2, 50× B27 without vitamin A, 1× glutamax, 1× NEAA, 1× antibiotic-antimycotic, 200 µM ascorbic acid, 100 µM cAMP, 1% ES-FBS, 10 µM DAPT, 20 ng/mL BDNF, 20 ng/mL GDNF, and the CEPT cocktail (50 nM Chroman 1 [HY-15392, MedChem Express], 5 µM emricasan (S7775, Selleckchem), 0.7 µM trans-ISRIB (#5284, Tocris), and polyamine supplement (#P8483, Sigma-Aldrich^116^) supplemented with 0.5% v/v GFR-Matrigel. The plate was left tilted and undisturbed in the incubator. After 3 days, 60% of the media in each well was replaced with fusion media. This was done slowly, while keeping the plate tilted with minimal disturbance to the “fused” organoid pair in each well. The same was done on D6 and D9. Subsequently, 80% of the media was replaced every 3 days. On D4, the plate was transferred to an orbital shaker at 80 rpm. The shaker speed was increased to 90 rpm on D5, 100 rpm on D6, and starting D7, the assembloids were kept at 110 rpm. Between 5 and 10 weeks postfusion, assembloids were harvested for electrophysiological experiments.

### Plasmids and lentiviruses

Synapsin-EGFP (hSyn-GFP) lentiviruses with the VSV-G pseudo-type were generated using the pHR-hSyn-eGFP plasmid^117^ (Addgene: 114215, a gift from Xue Han) by the St. Jude Viral Vector Core. For APEX2 experiments, pLenti-hSyn-V5-COX4-APEX2 plasmid was generated by cloning the V5-COX4-APEX2 sequence from pAAV-COX4-dAPEX2^65^ (Addgene: 117176, a gift from David Genty) into the pLenti backbone containing the human *SYN* promoter. Briefly, pAAV-COX4-APEX2 was digested with BspE1 and EcoR1, and pLenti-hSyn-nucGFP^118^ (Addgene: 140190, a gift from Lorenz Studer) was digested with EcoRI and AgeI to remove the nucGFP-coding sequence. Insert containing the V5-COX4-APEX2 sequence was then ligated into the pLenti-hSyn backbone. The resulting plasmid sequence was confirmed by Sanger sequencing. Lenti-hSyn-V5-COX4-APEX2 (hSyn-V5-Mito-APEX2) lentiviruses with the VSV-G pseudo-type were generated by the St. Jude Viral Vector Core.

Lentiviral vectors prepared at 1-3bn TU/mL were added to organoids in the bioreactor at 200×. For vectors at lower titer, 2 µg/mL Polybrene was also added to the media. After 18–20 h, organoids were washed twice with DMEM:F12 and fed with fresh media. Media changes were continued according to the protocol. Lentiviral expression was detected at 72 h posttransduction.

### Immunofluorescence and light microscopy

Organoids were briefly rinsed in phosphate-buffered saline (PBS) and then fixed in 4% paraformaldehyde in PBS overnight at 4°C. Following rinses in PBS, organoids were cryoprotected by incubation overnight in 30% sucrose in PBS. Organoids were then mounted in Optimal Cutting Temperature (O.C.T.) Compound (Tissue-Tek). Samples were stored at –80°C until cryosectioning. Cryosectioning was performed on a Leica CM 3050 cryostat set to –20°C. Serial sections of 20-µm thickness were mounted onto FisherBrand Superfrost Plus microscope slides and stored at –20°C.

Slides were briefly rehydrated with PBS and then blocked for 1 h at room temperature in blocking buffer (PBS, 5% normal donkey serum, 0.2% Triton-X100, 0.02% sodium azide, filter sterilized). Slides were incubated overnight at 4°C in primary antibodies diluted in blocking buffer, washed with PBS-Tween (0.1%), and incubated 1 h at room temperature in secondary antibodies diluted 1:500 in blocking buffer. Slides were then washed with PBS-Tween (0.1%), and nuclei were labeled with DAPI. Excess DAPI was removed by washing with PBS, and slides were dried and mounted for imaging using Prolong Diamond (Thermo Fisher, P36961). Images were acquired on a Zeiss Axio Imager M2 equipped with a 20× Plan-Apochromat Objective (Zeiss, 0.8 NA), 40× EC Plan-NeoFluar Objective (Zeiss, 1.3 NA), and Apotome.2 (Zeiss). Images including GABA were acquired using the 40× objective. All other images were acquired using the 20× objective. During imaging, exposure times were kept below saturation, and imaging conditions were constant within experiments. For images acquired with the Apotome.2, Z-series were acquired at software-recommended intervals and image stacks were then deconvolved using ZEN software and a constrained iterative algorithm. Images are shown as maximum intensity projections prepared in Zeiss ZEN 3.7 software.

The following primary antibodies and dilutions were used: TCF7L2 (Cell Signaling Technologies 2569, 1:1000), OTX2 (R&D Systems AF1979, 1:100), TUBB3 (Abcam Ab107216, 1µg/mL), SOX2 (R&D Systems MAB2018, 1:200), V5 (Invitrogen R960-25, 1:1000), FOXP2 (Abcam ab16046, 1:250), LHX2 (Sigma ABE1402,1:250), GBX2 (R&D Systems AF4638, 1:250), and GABA (Sigma A2052, 1:5000). The following f(ab′)_2_ secondary antibodies from Jackson Immunoresearch were used: donkey anti–rabbit Alexa Fluor (AF) 488 (711-546-152), donkey anti–goat AF 647 (705-606-147), donkey anti–mouse AF 488 (715-546-150), donkey anti–mouse AF 647 (715-606-150), and donkey anti–chicken AF 647 (703-606-155).

### Bulk RNA-seq

Each sample consisted of 2-3 pooled organoids from the indicated condition. Total RNA was isolated using the Direct-zol RNA Microprep Kits (Zymo, R2061), and DNA contamination was removed using the DNA-free DNA Removal Kit (Thermo Fisher, AM1906). RNA was quantified using the Quant-iT RiboGreen RNA assay (ThermoFisher) and quality checked by the 2100 Bioanalyzer RNA 6000 Nano assay (Agilent) or 4200 TapeStation High Sensitivity RNA ScreenTape assay (Agilent) prior to library generation. Libraries were prepared from total RNA with the TruSeq Stranded mRNA Library Prep Kit, according to the manufacturer’s instructions (Illumina PN 20020595). Libraries were analyzed for insert-size distribution using the 2100 BioAnalyzer High Sensitivity kit (Agilent), 4200 TapeStation D1000 ScreenTape assay (Agilent), or 5300 Fragment Analyzer NGS fragment kit (Agilent). Libraries were quantified using the Quant-iT PicoGreen ds DNA assay (ThermoFisher) or by low-pass sequencing with a MiSeq nano kit (Illumina). Paired-end 100-cycle sequencing was performed on a NovaSeq 6000 (Illumina) in the St. Jude Hartwell Center Genome Sequencing Core.

For tdTomato-expression analysis, we built a custom reference genome by adding the tdTomato sequence to the human genome (hg38, gencode v31). The tdTomato sequence was also added to the gene-annotation gtf file (gencode v31). Read alignment to the custom genome was performed with STAR (version 2.7) software.^119^ Gene-level read count was determined using RSEM (version 1.3.1).^120^

For differential gene expression analysis, only protein-coding genes validated at GENECODE confidence level 1 to 3 were considered. To remove genes that were lowly expressed, we first calculated the cutoff as 10 read counts per million library size, where the library size was defined as the median library size in the data set. We then kept genes with expression level (counts per million) equal to or above the cutoff in a minimum number of samples, where the number of samples was chosen according to the minimum group sample size. The data were then normalized by TMM function in edgeR package,^121^ followed by the limma package with its voom method, linear modeling, and empirical Bayes moderation to assess differential expression.^122^

Markers of interest were identified by performing a differential-expression analysis using *BrainSpan* Developmental Transcriptome data. Thalamic structures (mediodorsal nucleus of the thalamus and dorsal thalamus) were compared with all cortical structures. The top 100 up- or down-regulated genes in thalamic vs cortical structures were identified as “thalamic” or “cortical” markers, respectively.

Deconvolution of bulk RNA-seq data was performed using the VoxHunt (v1.0.1)^123^ R package using the default workflow (https://quadbio.github.io/VoxHunt/articles/deconvolution.html). *Allen Brain Atlas* data derived from E13 mouse brain were used as a reference. The “broad” gene set contained the top 50 markers for each region of interest. The top 15 markers were then used as input for the deconvolution tool.

GO term enrichment analysis was performed using g:Profiler.^124^ For all analyses, a custom background was uploaded containing genes detected in the data set of interest. Driver terms containing fewer than 300 genes were selected for graphing. All graphs, except heatmaps, were prepared in R using ggplot2 (v3.4.0).^125^ Heatmaps were prepared using the ComplexHeatmap (v2.10.0) R package.^126,127^

### RT-qPCR

RNA was isolated and DNase-treated, as described above. Reverse transcription was performed using 100 ng RNA and the iScript cDNA Synthesis Kit (Bio-Rad, 1708891). A qPCR analysis was then performed using SYBR Green Master Mix (Thermo Fisher, 4309155) and a C1000 Touch Thermal Cycler (Bio-Rad). The following primers were used: *GAPDH* Forward 5′-AATCCCATCACCATCTTCCA-3′, *GAPDH* Reverse 5′-TGGACTCCACGACGTACTCA-3′, *TCF7L2* Forward 5′-GAATCGTCCCAGAGTGATGTCG-3′, *TCF7L2* Reverse 5′-TGCACTCAGCTACGACCTTTGC-3′, *OLIG3* Forward 5′-TGAGGCTGAAGATCAACGGACG-3′, *OLIG3* Reverse 5′-AGTTTCTGGCGAGCAGGAGTGT-3′, *GBX2* Forward 5′-GCGGAGGACGGCAAAGGCTTC-3′, *GBX2* Reverse 5′-GTCGTCTTCCACCTTTGACTCG-3′, *LHX9* Forward 5′-ACCTGCTTTGCCAAGGACGGTA-3′, *LHX9* Reverse 5′-TGACCATCTCCGAGGCGGAAAT-3′, *OTX2* Forward 5′-GGAAGCACTGTTTGCCAAGACC-3′, *OTX2* Reverse 5′-CTGTTGTTGGCGGCACTTAGCT-3′, *FOXG1* Forward 5′-GTATGTGGTCACTAACAGGTC-3′, and *FOXG1* Reverse 5′-ACCACAGTATCACAATCAAG-3′.

Data were analyzed using the 2^−ΔΔCq^ method [previously known as the 2^−ΔΔCt^ method, first described in the Applied Biosystems User Bulletin 2 (P/N 4303859)].^128^ Transcripts of interest were normalized first to *GAPDH* (within sample), then to the mean ΔCq of the hThO samples with high tdTomato that were previously used for bulk RNA-seq. Regression analyses were performed in R using normalized values and graphed using the ggscatter function from ggpubr (v0.5.0). For all transcripts, except *FOXG1*, r and p were calculated using the Pearson correlation method, where *r* represents the correlation coefficient and p represents the p-value, and lines were fit using linear regression. Due to the presence of extreme outliers, for *FOXG1* the *r*_s_ and p-value were calculated using the Spearman correlation method, in which *r*_s_ represents the correlation coefficient, p represents the p-value, and the nonlinear curve is fit using the Loess local polynomial-regression method.

### Preparation and sequencing of the snRNA-seq library

Two independent biological replicates were performed per the differentiation protocol (either cortical or thalamic), each containing 36 organoids pooled together. The hThOs were D91 or D96; the hCOs were D91. All organoids were flash frozen in liquid nitrogen and stored at –80°C until dissociation. Nuclei dissociation was performed as previously described.^129^ Briefly, frozen tissue was mechanically dissociated with a Dounce homogenizer in detergent lysis buffer containing 0.32 M sucrose, 10 mM HEPES (pH 8.0), 5 mM CaCl_2_, 3 mM magnesium acetate, 0.1 mM EDTA, 1 mM DTT, and 0.1% Triton-X100. The resulting homogenate was filtered through a 40-µm strainer and washed with the same solution described, without the Triton-X100 added. Nuclei were then centrifuged at 3200 ×*g* for 10 min at 4°C, and the supernatant was decanted. A sucrose-dense solution containing 1 M sucrose, 10 mM HEPES (pH 8.0), 3 mM magnesium acetate, and 1 mM DTT was carefully layered underneath the remaining supernatant and then spun at 3200 ×*g* for 20 min at 4°C. The supernatant was discarded, and the final remaining nuclei were resuspended in 0.4 mg/mL BSA and 0.2 U/µL Lucigen RNAse inhibitor (catalog number 30281-1) diluted in PBS. Between 5000 and 10,000 nuclei were inspected and counted on a hemacytometer before loading onto the 10× Chromium Controller (10× Genomics, catalog number 1000171). The snRNA-seq libraries were prepared using the 10× Genomics Chromium Next GEM Single Cell Kit, version 3.1 single index gene expression profiling assay, according to the manufacturer’s instructions.

Libraries were analyzed for insert-size distribution by using the 2100 BioAnalyzer High Sensitivity kit (Agilent), 4200 TapeStation D1000 ScreenTape assay (Agilent), or 5300 Fragment Analyzer NGS fragment kit (Agilent). Libraries were quantified using the Quant-iT PicoGreen ds DNA assay (ThermoFisher) or by low-pass sequencing with a MiSeq nano kit (Illumina). Paired-end 100-cycle sequencing was performed on a NovaSeq 6000 (Illumina) in the St. Jude Hartwell Center Genome Sequencing Core.

### Analysis of snRNA-seq data

Sequences from each Illumina-sequencing data set were de-multiplexed using bcl2fastq v2.20.0.422 (Illumina). The sequencing data were aligned to the human reference genome GRCh38 (10× Genomics, v2020-A) using the CellRanger “count” algorithm (10× Genomics, v7.0.0); however, the “–force-cells” option was set to the estimated number of cells loaded for each sample (snCBO1: 6,000; snCBO2: 8,000; snTha1: 8,000; snTha2: 10,000). From the gene expression matrix, the downstream analysis was carried out in R (v4.2.1). First, the ambient RNA signal was removed using the default SoupX (v1.6.2) workflow (autoEstCounts and adjustCounts; github.com/constantAmateur/SoupX).^130^

Each data set was initially filtered so that genes that were expressed in at least three cells, and cells that expressed at least 200 genes were included. Additionally, cells with fewer than 300 genes (presumed to be droplets or cellular debris), fewer than 500 UMIs, more than 1% unique transcripts derived from mitochondrial genes, or more than 3 median absolute deviations (MADs) from the median number of unique transcripts derived from mitochondrial genes were removed. Afterwards, cells with more than 3 MADs from the median number of genes expressed were removed.

Samples were then preprocessed using the standard Seurat (v4.3.0) workflow (NormalizeData, ScaleData, FindVariables, RunPCA, FindNeighbors, FindClusters, and RunUMAP; github.com/satijalab/Seurat).^131–134^ Data sets were individually log-normalized using Seurat’s NormalizeData with default parameters. Cell cycle scoring was conducted using the associated S- and G2M-phase gene list from Tirosh et al.^135^ and the CellCycleScoring command in Seurat. We calculated 3000 features exhibiting high cell-to-cell variation in the data set by using Seurat’s FindVariableFeature function. Next, we scaled the data by linear regression against the number of reads by using Seurat’s ScaleData function with default parameters. The variable genes were projected onto a low-dimensional subspace by performing principal component analysis using Seurat’s RunPCA function with default parameters. The number of principal components (n = 30) was selected based on inspection of the plot of variance explained.

Data sets were integrated using Harmony (v 0.1.1) with default parameters.^136^ A shared-nearest-neighbor graph was constructed based on the Euclidean distance in the low-dimensional subspace using Seurat’s FindNeighbors with dims = 1:30 and default parameters. Integrated data sets then underwent nonlinear dimensional reduction and visualization using UMAP. Clusters were identified using a resolution of 0.4 and the Leiden algorithm for the integrated data sets. Pseudotime analysis was conducted using Monocle3 (v1.3.1) with default parameters.^137–140^ Trajectory starting points were manually selected based on the expression of mitotic markers (e.g., *MKI67*) and/or neural precursor markers (e.g., *TNC*). Mapping of snRNA-seq data sets onto *Allen Brain Atlas* and *BrainSpan* reference data sets was performed using the VoxHunt (1.0.1) R package with the suggested workflows (https://quadbio.github.io/VoxHunt/articles/getting_started.html; https://quadbio.github.io/VoxHunt/articles/other_references.html).^123^ For *Allen Brain Atlas* comparisons, data derived from E15 mouse embryos were used. For *BrainSpan* comparisons, data derived from human fetal tissue 13–24 postconception weeks (pcw) were used. Neural communication patterns were predicted and visualized using the NeuronChat (v1.0.0) R package with the suggested workflow (https://github.com/Wei-BioMath/NeuronChat/blob/main/vignettes/NeuronChat-Tutorial.html).^58^

Cell types were assigned to each cell/cluster based on marker expression and cell cycle analysis. For hThO annotation, markers of interest were identified based on a comparison to previously published scRNA-seq or snRNA-seq studies in developing mouse thalamus or diencephalon.^141,142^ Additional markers were identified based on previously published *in situ* studies examining embryonic rodent thalamus. For example, within the mouse thalamus^143^ and hThOs, *SOX2* is expressed in a subset of postmitotic neurons. For hCO cluster annotation, markers of interest were identified using a previously published scRNA-seq study that examined human neocortex at midgestation.^56^ Additional markers were identified based on previously published *in situ* studies examining embryonic rodent cortex.

### Electron microscopy for DAB-labeled samples

Prior to fusion, TCF7L2-tdTomato^+^ thalamic and VGLUT1-tdTomato^+^ hSyn-GFP^+^ cortical organoids were separately transduced in low-attachment 6-well plates with 10^7^ TU/mL Lenti-hSyn-V5-COX4-APEX2 lentiviral vector. After 18–20 h, organoids were washed twice with DMEM:F12 and fed fresh media. After 3 days, APEX2-transduced thalamic organoids were fused with cortical organoids, and APEX2-transduced cortical organoids were fused with thalamic organoids to generate the assembloids described above. At 6–7 weeks postfusion, the assembloids were prepared for electron microscopy analysis. Specifically, each assembloid was embedded at the center of a UV-sterilized Nunc Thermanox plastic coverslip (Thermo Fisher, 174950) in 5 µL GFR-Matrigel for 1 h at 37°C. They were then transferred to the fusion media in 6-well plates and placed in the incubator overnight. The following day, a sterile blade was used to cut a V-shaped notch out of the coverslip on the side containing the APEX2^+^ half of the assembloid. DAB labeling was then performed using an adapted protocol.^144^

Briefly, after 1 additional day at 37°C, assembloids were fixed for 1 h in 2% glutaraldehyde in 0.1 M sodium cacodylate at room temperature, after which the fixative was replaced, and samples were incubated 1 h at 4°C. The samples were then washed thrice for 5 min in ice-cold wash solution (0.1 M sodium cacodylate). Next, the samples were incubated 5 min in 20 mM glycine in 1× sodium cacodylate, then washed thrice for 5 min on ice. The samples were preincubated in 0.5 mg/mL DAB in 0.1 M sodium cacodylate for 30 min on ice. The samples then underwent DAB staining in 0.5 mg/mL DAB and 50 mM H_2_O_2_ in 0.1 M sodium cacodylate on ice. The reaction was terminated by washing the samples thrice for 5 min on ice in wash solution.

After the DAB labeling developed, samples were postfixed in 2% osmium tetroxide in 0.1 M cacodylate buffer on ice for 30 min. Samples were subsequently washed 5 times for 2 min in ice-cold water, and then they were contrasted with 2% uranyl acetate overnight at 4°C. Samples were washed five times for 2 min in ice-cold water. Samples were dehydrated on ice by an ascending series of ethanol to 100%, followed by 100% propylene oxide at room temperature. Samples were infiltrated with EmBed-812 and polymerized at 60°C. Embedded samples were sectioned at ∼70 nm on a Leica ultramicrotome and examined in a ThermoFisher Scientific TF20 transmission electron microscope at 80 kV. Digital micrographs were captured with an Advanced Microscopy Techniques imaging system. Unless otherwise indicated, all reagents were from Electron Microscopy Sciences.

### Identification of synapses by transmission electron microscopy

Individual organoids were harvested between D102 and D121 for electron microscopy imaging. Samples were fixed in 0.1 M cacodylate buffer containing 2.5% glutaraldehyde and 2% paraformaldehyde. Samples were postfixed in reduced osmium tetroxide and contrasted with aqueous uranyl acetate. Dehydration was by an ascending series of ethanol to 100%, followed by 100% propylene oxide. Samples were infiltrated with EmBed-812 and polymerized at 60°C. Embedded samples were sectioned at ∼70 nm on a Leica ultramicrotome and examined in a ThermoFisher Scientific TF20 transmission electron microscope at 80 kV. Digital micrographs were captured with an Advanced Microscopy Techniques imaging system. Unless otherwise indicated, all reagents were from Electron Microscopy Sciences.

### Fusion of 2-dimensional organoids

TCF7L2-tdTomato^+^ hThOs and VGLUT1-tdTomato^+^ hSyn-GFP^+^ hCOs were halved using a pair of ultra-fine clipper scissors and plated in a culture-insert 2-well in μ-dish 35 mm (81176, Ibidi). Specifically, each half chamber was first coated with human ES-qualified Matrigel diluted 1:200 in DMEM:F12, for 1 h at room temperature. The coating solution was aspirated and 100 µL fusion media was added to each half. One hThO half was placed in one chamber and 1-2 hCO halves were placed in the other and allowed to attach and extend neural processes for 5 days. On D5, the barrier was removed using sterilized blunt forceps, and the organoids were maintained in culture with media changes every 7 days. The barrier region in each dish was imaged every 3–7 days, from D9 to D61, at the same exposure time on a Zeiss AxioObserver D1.

### Whole-cell patch-clamp electrophysiology

Whole-cell recordings were made in individual organoids between D90 and D141 or in assembloids between D19 and D78 postfusion. Organoids were placed in a recording chamber mounted on a two-photon laser-scanning microscope (Bruker) and superfused (2-3 mL/min) with aCSF containing the following solution: 125 mM NaCl, 2.5 KCl, 2 mM CaCl_2_, 2 mM MgCl_2_, 1.25 mM NaH_2_PO_4_, 26 mM NaHCO_3_, and 20 mM glucose at 300–310 mOsm, equilibrated with 95% O_2_/5% CO_2_ at 32°C.

Cells were visualized under two-photon guidance by using PrairieView v5.5 software. Whole-cell voltage- and current-clamp recordings were made from visually identified thalamic or cortical cells. Short-term synaptic plasticity, 1-Hz induction of LTD, and 40-Hz LTP were recorded in voltage-clamp mode (V_Hold_ = –60 mV), with an internal pipette containing the following solution: 125 mM CsMeSO_3_, 2 mM CsCl, 10 mM HEPES, 0.1 mM EGTA, 4 mM ATP-Mg_2_, 0.3 mM GTP-Na, 10 mM creatine phosphate-Na_2_, 5 mM QX-314 chloride, and 5 mM TEA-Cl at pH 7.4 and 290-295 mOsm. Borosilicate glass pipettes (Sutter, 3-6 MΩ open pipette resistance) were used.

For investigating membrane properties and LTP induced by the STDP protocol, the internal solution contained 115 mM potassium gluconate (KGlu), 20 mM KCl, 10 mM HEPES, 4 mM MgCl_2_, 0.1 mM EGTA, 4 mM ATP-Mg_2_, 0.4 mM GTP-Na, and 10 mM creatine phosphate-Na_2_ at pH 7.4 and 290-295 mOsm. Recordings were obtained using a Multiclamp 700B amplifier (Axon Instruments). Signals were digitized with an Axon Digidata 1550B (Axon Instruments) at 20 kHz and filtered at 2 kHz using Clampex 10.7 software. The liquid-junction potential was calculated to be –10 mV and was corrected for in each recording.

In current-clamp experiments, the rheobase was measured by first injecting a hyperpolarizing current step (–20 pA), followed by a depolarizing ramp (from –20 pA to +200 pA) into cells. The current at which the first AP was generated was recorded and averaged across cells. A series of hyperpolarizing and depolarizing step currents were injected into cells in current-clamp mode (+10 pA steps from –50 pA to +240 pA for 1 s) to measure input resistance and evoked firing rates.

In voltage-clamp experiments, synaptic currents were evoked using a bipolar concentric stimulating electrode (World Precision Instruments) or a homemade 2-prong stimulating electrode connected to a stimulus-isolation unit (Iso-Flex, A.M.P.I.) positioned in either the thalamic or cortical side of assembloids. Stimulus intensities were adjusted prior to each experiment to elicit measurable EPSCs in cortical or thalamic neurons. Because of the variability between assembloids, the amplitudes of evoked EPSCs ranged from –20 pA to –300 pA. PPRs were measured by delivering 2 stimuli 50, 100, 200, 500, or 1000 ms apart for both TC and CT synapses.

TC LTP was induced by high-frequency stimulation: 10 trains of 40-Hz stimulation for 200 ms every 5 s, repeated 3× every 5 min.^91^ Additional TC LTP and CT LTP were induced via an STDP protocol: a single presynaptic electrical stimulation preceded four postsynaptic APs by 10 s. Postsynaptic APs were induced by four somatic current injections of 2 nA (2-ms duration) at 40 Hz. This protocol was repeated 50 times (at 1 Hz) every 5 min, for a total of three times (long STDP-induction protocol).^145^ In a subset of experiments, the STDP protocol was delivered only 1× (short STDP-induction protocol). LTD was induced at TC and CT synapses by delivering electrical low-frequency stimulation at 1 Hz for 900 pulses.^146^ In all long-term synaptic plasticity induction protocols, cells were current-clamped at –60 mV. EPSC peak amplitudes were measured before and after synaptic plasticity induction in voltage-clamp mode (V_hold_ = –60 mV) using paired electrical stimulation (10 Hz) delivered every 20 s for a 5-min baseline period and a 30-min postinduction period.

All electrophysiological experiments were analyzed offline using Clampfit 10.7 software. For all long-term synaptic plasticity experiments, raw EPSC amplitudes were measured, averaged per minute, and expressed as a percent change from baseline. The amplitude of the first EPSC peak was measured if a polysynaptic response was elicited. To determine a change in synaptic strength after the plasticity-induction protocols, the full postinduction time periods of all the cells in the experiment were averaged and compared to a theoretical baseline of 100% by using a one-sample *t*-test (GraphPad Prism 8.4.2), unless noted. To compare between experimental drug conditions, a one-way ANOVA with Dunnett’s multiple comparisons post-hoc test was used. PPR was calculated by measuring the peak amplitude of the evoked EPSC from both pulses and dividing the EPSC2 peak amplitude by the EPSC1 peak amplitude. The PPR for each ISI of each synapse was compared against 1.0 by using a one-sample *t*-test and between TC and CT synapses by using an unpaired two-tailed *t*-test.

### Two-photon calcium imaging

Two-photon calcium imaging was performed as described previously.^73^ Briefly, two-photon laser-scanning microscopy was performed using an Ultima imaging system (Bruker), a Ti:sapphire Chameleon Ultra femtosecond-pulsed laser (Coherent, 820 nm) and 60× [0.9 numerical aperture] water-immersion infrared objectives (Olympus). Fluo-5F (300 μM) and Alexa 594 (10-25 μM) were included in the internal solution containing 115 mM potassium gluconate, 20 mM KCl, 10 mM HEPES, 4 mM MgCl_2_, 0.1 mM EGTA, 4 mM ATP-Mg_2_, 0.4 mM GTP-Na, 5 mM QX-314 chloride, and 10 mM creatine phosphate-Na_2_ at pH 7.4 and 290-295 mOsm. Synaptically evoked changes in fluorescence of both fluorophores were measured in line-scan mode in a dendritic spine and the parent dendritic shaft. Line scans were analyzed as a ratio of normalized green (G) (Fluo-5F) fluorescence to normalized red (R) (Alexa Fluor 594) fluorescence (G/R). A line-scan was performed through every visible dendritic spine on a targeted dendritic branch, in an orientation that was parallel to the dendritic spine neck and orthogonal to the dendritic shaft.

### Statistical analyses

Statistical tests were performed using Prism (Graphpad) or Sigmaplot (Systat) software. Statistical comparisons are noted in the text or figure legends. Unless otherwise noted, distributions were tested for normality (Shapiro-Wilk test) and equal variance (Brown-Forsythe test). If the distribution passed, a paired or unpaired *t*-test was performed. If it failed, a rank-sum test or signed-rank test was performed. To compare more than two distributions, a one-way or repeated-measures ANOVAs was performed. Significance was designated as *P* <0.05. All data are presented as the mean ± SEM, and the sample size (N) is presented as the number of cells per the number of assembloids.

### Drugs

All salts for aCSF were purchased from Sigma-Aldrich. QX-314 chloride was purchased from Hello Bio. DL-AP5 and MPEP were purchased from Tocris Bioscience. To create stock solutions, MPEP was dissolved in DMSO and DL-AP5 was dissolved in water; both were kept frozen at – 20°C until dilution in aCSF to the final concentration. For iBAPTA experiments, BAPTA tetra-potassium salt and BAPTA tetra-cesium salt were included in KGlu- and Cs-based intracellular solutions, respectively, at 20 mM.

