## Supplemental Data for "Synaptic plasticity in human thalamocortical assembloids"

### Supplemental Figures

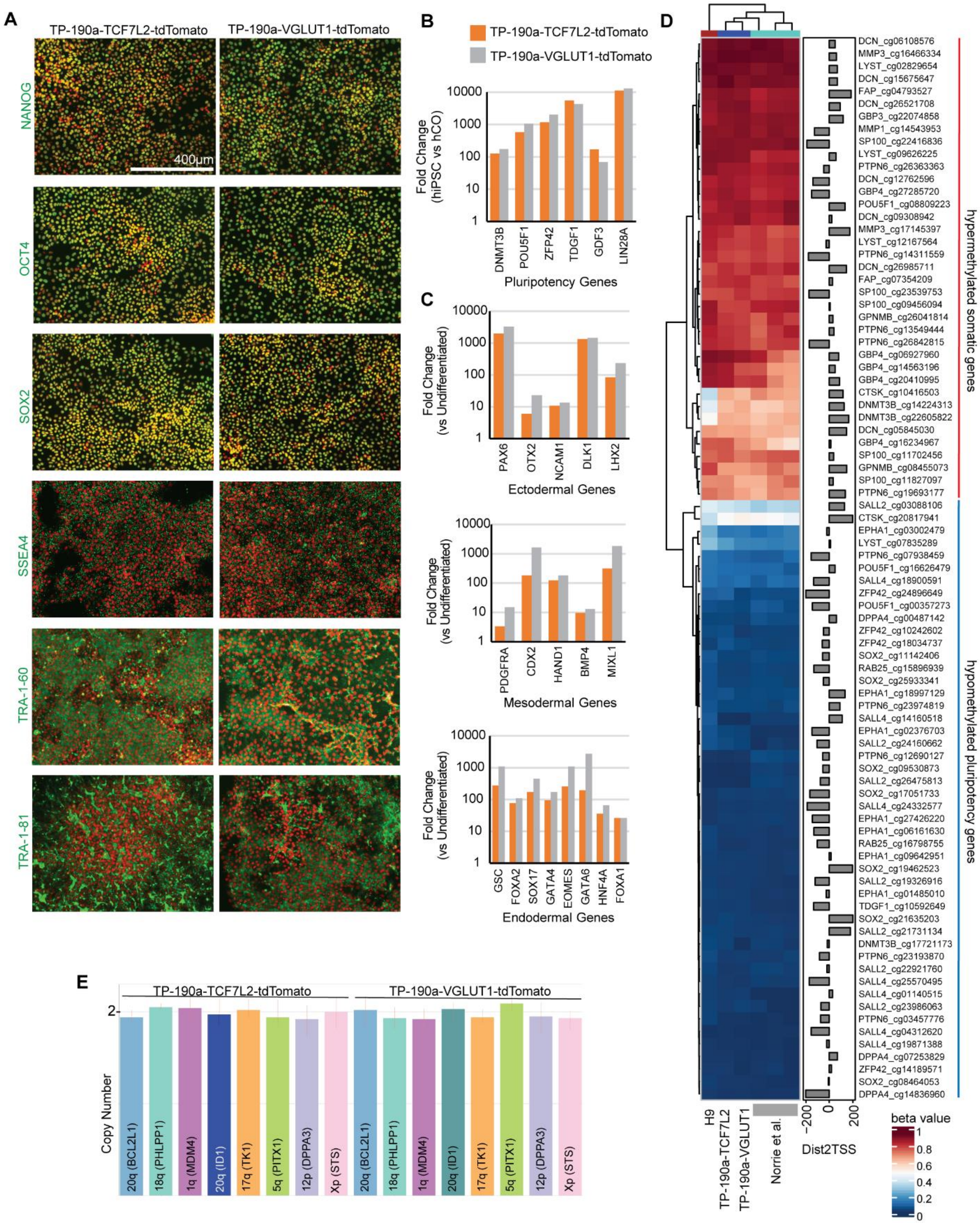


**Figure S1. Validation of TP-190a reporter lines (related to Figure 1).**

(**A**) Immunostaining of undifferentiated TP-190a-TCF7L2-tdTomato and TP-190a-VGLUT1-tdTomato colonies shows expression of pluripotency markers NANOG, OCT4, SOX2, SSEA4, TRA-1-60, and TRA-1-81. DAPI is shown in red, and the pluripotency markers are shown in green.
(**B**) RT-qPCR of undifferentiated TP-190a-TCF7L2-tdTomato and TP-190a-VGLUT1-tdTomato colonies shows expression of pluripotency genes *DNMT3B*, *POU5F1* (*NANOG*), *ZFP42*, *TDGF1*, *GDF3*, and *LIN28A*. Fold change in gene expression is shown compared to that in 13-week-old hCOs and on a logarithmic scale.
(**C**) RT-qPCR of lineage-specific markers in TP-190a-TCF7L2-tdTomato and TP-190a-VGLUT1-tdTomato lines differentiated into ectodermal, mesodermal, and endodermal lineages. Fold change in gene expression is shown compared to that of undifferentiated iPSCs and on a logarithmic scale.
(**D**) DNA-methylation status at select epigenetic markers in TP-190a-TCF7L2-tdTomato and TP-190a-VGLUT1-tdTomato lines shows hypermethylation of somatic genes and hypomethylation of pluripotency genes, similar to findings in established, published hPSC lines.
(**E**) RT-qPCR–based copy number variation assay in TP-190a-TCF7L2-tdTomato and TP-190a-VGLUT1-tdTomato lines shows normal copy number in seven regions of the genome that together carry more than 90% of recurrent copy number variations found in hiPSCs and human embryonic stem cells (i.e., 20q, 18q, 1q, 17q, 5q, 12p, and Xp).

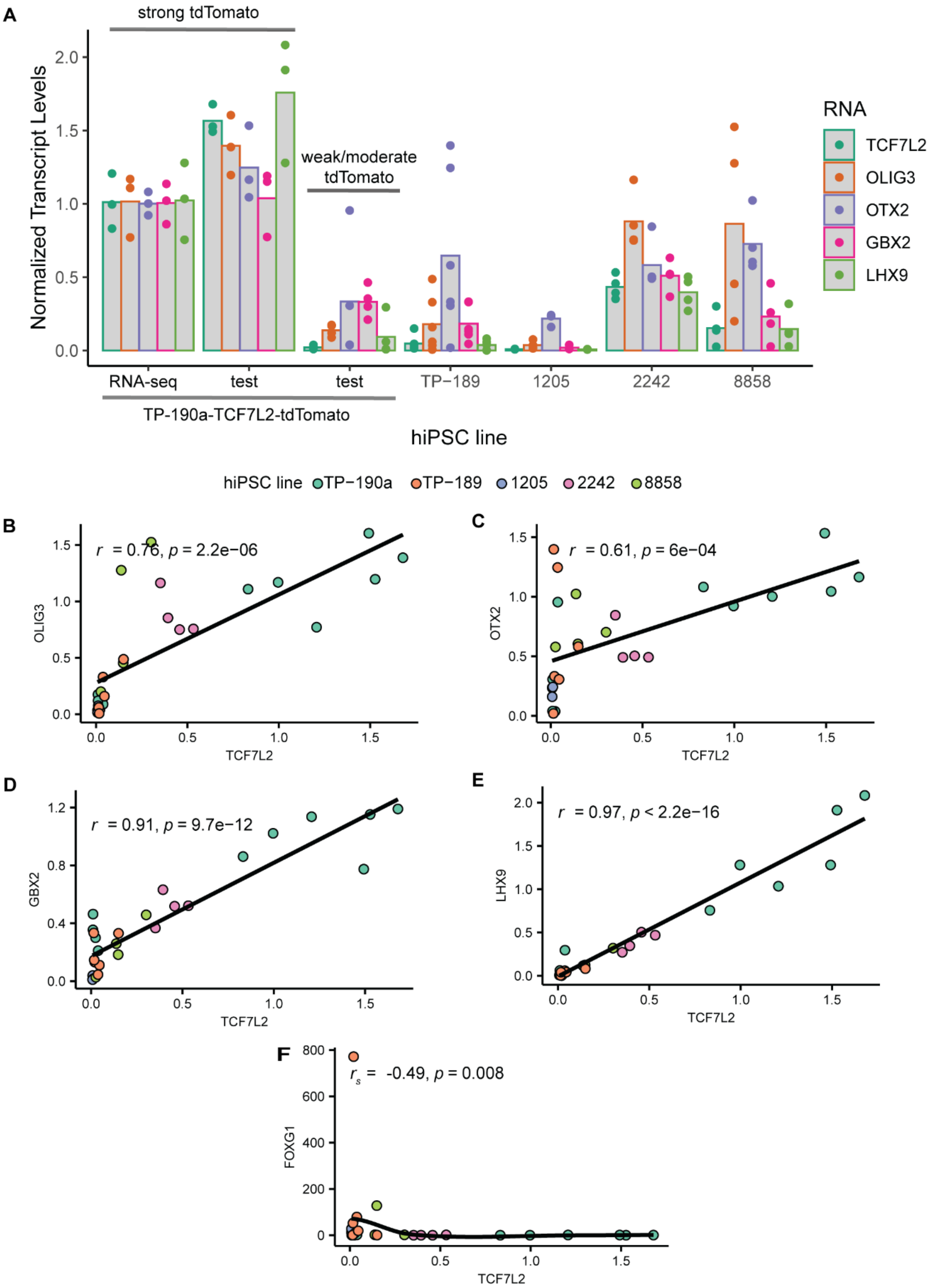


**Figure S2. *TCF7L2* predicts thalamic marker expression in hThOs derived from multiple hiPSC lines (related to Figure 1).**

(**A**) Bar graph summarizing the expression of thalamic marker transcripts, as detected by RT-qPCR in hThOs from multiple hiPSC lines. Each point represents a sample containing two to three pooled organoids. “RNA-seq” indicates that the sample was previously used for RNA-seq and confirmed to have strong thalamic marker expression. All other samples were used for RT-qPCR only. “Strong tdTomato” or “weak/moderate tdTomato” indicates tdTomato fluorescence, as assessed visually before RNA isolation. The 2242 and 8858 lines successfully generated organoids with moderate expression of *TCF7L2* and other thalamic markers.
(**B**) *TCF7L2* positively predicts the level of *OLIG3*, a marker of neural precursors in the developing thalamus.
(**C**) *TCF7L2* also positively predicts the levels of *OTX2*, another marker of neural precursors in the developing thalamus.
(**D**) *TCF7L2* positively predicts the level of the thalamic neuron marker *GBX2*.
(**E**) *TCF7L2* positively predicts the level of *LHX9*, another thalamic neuron marker.
(**F**) *TCF7L2* is negatively correlated with the cortical marker *FOXG1*.

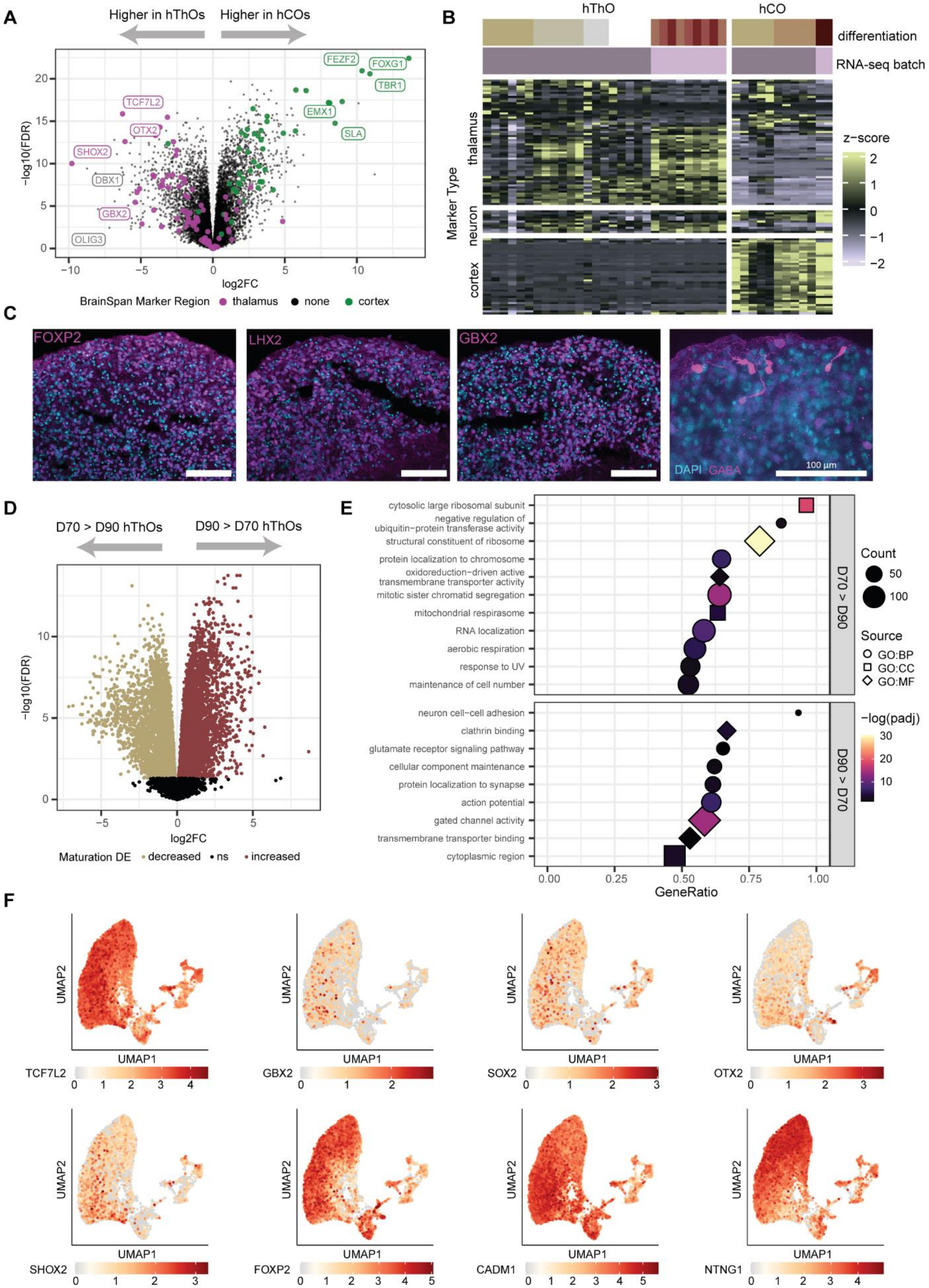
A
Higher in hThOs
Higher in hCOs
FEZF2
FOXG1
TBR1
TCF7L2
EMX1
OTX2
SLA
SHOX2
DBX1
GBX2
OLIG3
-log10(FDR)
log2FC
BrainSpan Marker Region
thalamus
none
cortex
B
hThO
hCO
differentiation
RNA-seq batch
Marker Type
thalamus
neuron
cortex
z-score
C
FOXP2
LHX2
GBX2
DAPI
GABA
100 µm
D
D70 > D90 hThOs
D90 > D70 hThOs
-log10(FDR)
log2FC
Maturation DE
decreased
ns
increased
E
cytosolic large ribosomal subunit
negative regulation of ubiquitin-protein transferase activity
structural constituent of ribosome
protein localization to chromosome
oxidoreduction-driven active transmembrane transporter activity
mitotic sister chromatid segregation
mitochondrial respirasome
RNA localization
aerobic respiration
response to UV
maintenance of cell number
neuron cell-cell adhesion
clathrin binding
glutamate receptor signaling pathway
cellular component maintenance
protein localization to synapse
action potential
gated channel activity
transmembrane transporter binding
cytoplasmic region
D70 > D90
D90 > D70
GeneRatio
Count
Source
GO:BP
GO:CC
GO:MF
-log(padj)
F
UMAP1
UMAP2
TCF7L2
GBX2
SOX2
OTX2
SHOX2
FOXP2
CADM1
NTNG1

**Figure S3. Validation of hThOs (related to Figure 1).**
All organoids were generated using TCF7L2 (hThO) or VGLUT1 (hCO) reporter lines and expressed strong tdTomato fluorescence, as determined by visual assessment.
(**A**) Volcano plot comparing Day (D) 69–D70 hThOs to D69–D72 hCOs. $Log_2$-fold change (FC) >0 indicates that a transcript is expressed at higher levels in hCOs than in hThOs, and $Log_2$FC <0 indicates that a transcript is expressed at higher levels in hThOs. The hThOs express significantly higher levels of many thalamic markers identified using *BrainSpan* (e.g., *TCF7L2*, *GBX2*, and *OTX2*) and those identified in the literature (e.g., *OLIG3*). Conversely, hCOs express significantly higher levels of *BrainSpan* cortical markers (e.g., *FOXG1*, *TBR1*, and *FEZF2*).
(**B**) Heatmap comparing D69–D70 hThOs to D69–D72 hCOs. Thalamus and cortex markers were identified using *BrainSpan*. Differentiations represent unique hiPSC differentiation experiments. The heatmap contains the following neuronal markers: *DCX*, *TUBB3*, *NEUROD1*, *RBFOX3*, *MAP2*, *NEFL*, *SYP*, *DLG4*, *CALB1*, *GAP43*, and *NCAM1*. The hThOs from multiple differentiations and sequencing batches express higher levels of thalamic markers, whereas the hCOs express higher levels of cortical markers. No clear patterns of neuronal markers emerge.
(**C**) Immunofluorescence images of D92 hThOs. Most cells express markers consistent with glutamatergic neurons in the thalamus: FOXP2, LHX2, and GBX2. GABA was detected in a subset of cells. Scale bars: 100 μm.
(**D**) Volcano plot comparing D69–D70 hThOs to D91–D94 hThOs. Color represents differential expression with maturation. Increased (D91–D94 > D69–D70): false discovery rate (FDR) <0.05 and $log_2$FC >0. Decreased (D69–D70 > D91–D94): FDR <0.05 and $log_2$FC <0.
(**E**) GO-term enrichment analysis of transcripts differentially expressed during hThO maturation. BP: biological process, CC: cellular component, MF: molecular function.
(**F**) UMAP plots of thalamic markers *TCF7L2*, *GBX2*, *SOX2*, *OTX2*, *SHOX2*, *FOXP2*, *CADM1*, and *NTNG1.* Color indicates normalized transcript level. Data were generated by snRNA-seq of D90 hThOs.
For (**A**) and (**B**), n = 29 hThO and 12 hCO samples. For (**D**), n = 9 D69–D70 hThOs and 8 D91–D94 hThOs. For (**D**) and (**E**), analysis was limited to hThOs from the same sequencing batch.

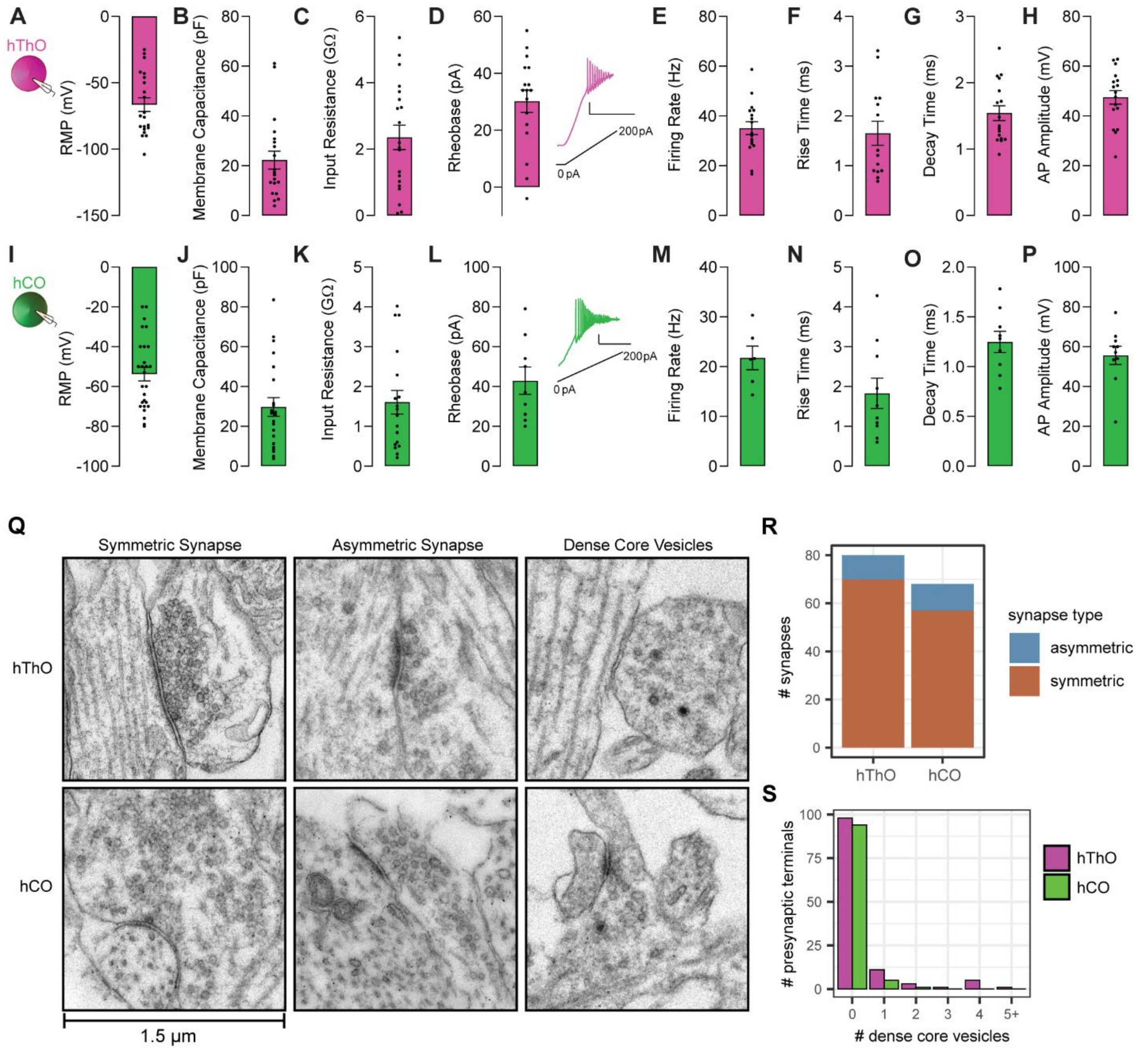

**Figure S4. Membrane and synapse properties of cells recorded in hThOs and hCOs (related to Figures 1 and 2).**

(**A**) Inset: Schematic of the recording configuration. Whole-cell patch-clamp recordings were made from cells in unfused hThOs. Bar graph of the average resting membrane potential (RMP) of hThO cells (–66.41 ± 5.12 mV, n = 21).

(**B**) Bar graph of the average membrane capacitance (Cm) of hThO cells (22.28 ± 3.65 pF, n = 20).

(**C**) Bar graph of the average input resistance (Rin) in response to a –50 pA hyperpolarizing step current injection into hThO cells (2356 ± 365.5 MΩ, n = 20).

(**D**) Left: Bar graph of the average minimum current injected into hThO cells to reach action potential (AP) threshold or rheobase (30.18 ± 3.94 pA, n = 17). Right: Example trace of an hThO cell responding to a depolarizing current injection ramp, from 0 to 200 pA. Scale bars: 20 mV, 1 s.

(**E**) Bar graph of the average firing rate in response to a depolarizing current ramp in hThO cells (35.13 ± 2.58 Hz, n = 17).

(**F**) Bar graph of the average AP rise time of hThO cells (1.65 ± 0.24 ms, n = 14).

(**G**) Bar graph of the average AP decay time of hThO cells (1.54 ± 0.11 ms, n = 17).
(**H**) Bar graph of the average AP amplitude of hThO cells (47.50 ± 2.72 mV, n = 17).
(**I**) Inset: Schematic of the experimental condition. Whole-cell patch-clamp recordings were made from cells in unfused hCOs. Bar graph of the average RMP of hCO cells (–53.74 ± 3.49 mV, n = 27).
(**J**) Bar graph of the average Cm of hCO cells (29.69 ± 4.67 pF, n = 22).
(**K**) Bar graph of the average Rin in response to a –50 pA hyperpolarizing step current injection into hCO cells (1606 ± 296.6 MΩ, n = 18).
(**L**) Left: Bar graph of the average minimum current injected to reach hCO AP threshold (42.89 ± 6.87 pA, n = 9). Right: Example trace of an hCO cell responding to a depolarizing current injection ramp, from 0 pA to +200 pA. Scale bars: 20 mV, 0.5 s.
(**M**) Bar graph of the average firing rate in response to a depolarizing current ramp in hCO cells (21.75 ± 2.36 Hz, n = 6).
(**N**) Bar graph of the average AP rise time of hCO cells (1.83 ± 0.38 ms, n = 10).
(**O**) Bar graph of the average AP decay time of hCO cells (1.25 ± 0.11 ms, n = 9).
(**P**) Bar graph of the average AP amplitude of hCO cells (55.61 ± 4.6 mV, n= 10).
(**Q**) Representative TEM images of symmetric synapses, asymmetric synapses, and presynaptic terminals containing dense core vesicles.
(**R**) Bar graph representing the number of symmetric and asymmetric synapses observed in TEM images from hThOs and hCOs. Chi square test, p >0.05.
(**S**) Histogram representing the number of dense core vesicles per presynaptic terminal observed in TEM images from hThOs and hCOs. Chi square test, p >0.05.
(**A**) to (**P**) Data shown are mean ± SEM with individual data points overlaid as dots.

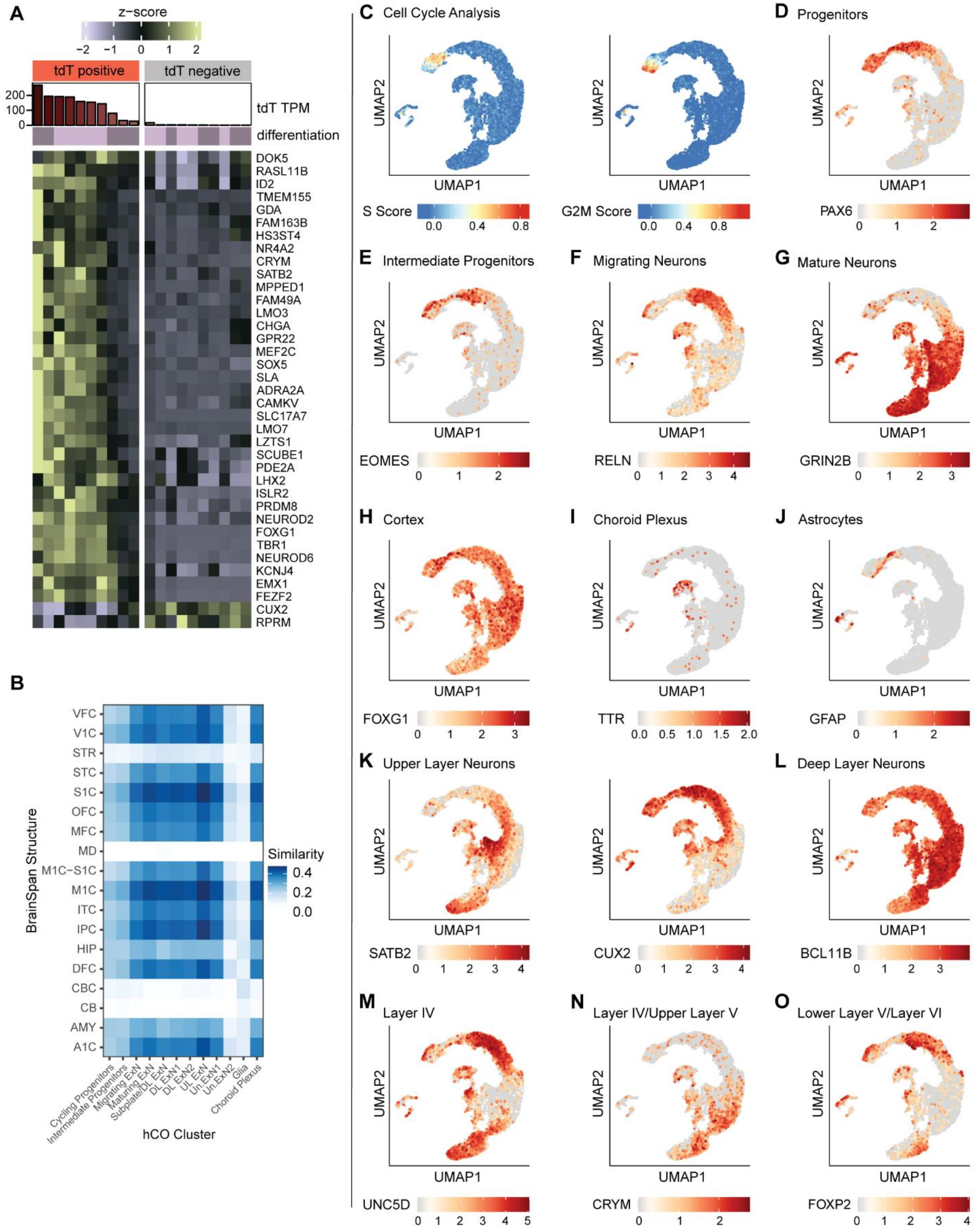


**Figure S5. Validation of hCOs (related to Figure 2).**

All hCOs were generated from the TP-190a-VGLUT1-tdTomato reporter line.

(**A**) Heatmap showing bulk RNA-seq data from D69–D72 hCOs. Organoids were visually categorized as positive or negative for tdTomato (tdT) fluorescence prior to sequencing. High *tdTomato* RNA levels (shown in tdT TPM) predict high expression of cortical markers identified using *BrainSpan*. Notably, the expression of *FOXG1*, *FEZF2*, and *TBR1* were higher in dtT$^+$ organoids than in dtT$^-$ organoids.
(**B-O**) The snRNA-seq analysis of D90 hCOs. All data were generated using VGLUT1-tdTomato$^+$ hCOs.
(**B**) VoxHunt analysis by cell cluster. Most clusters exhibit the highest correlations with *BrainSpan* samples from human neocortex (VFC, V1C, STC, S1C, OFC, MFC, M1C-S1C, M1C, ITC, IPC, DFC, A1C), aged 13–24 postconception weeks (pcw).
(**C**) UMAP plots showing cell cycle analysis results. Most cells undergoing mitosis are found within the Cycling Progenitor cluster. Color indicates S Score (left) or G2M Score (right).
(**D**) UMAP plot of the forebrain progenitor marker *PAX6*.
(**E**) UMAP plot of the intermediate progenitor marker *EOMES* (*TBR2*).
(**F**) UMAP plot of the neuronal migration marker *RELN*.
(**G**) UMAP plot of the NMDAR subunit and mature neuron marker *GRIN2B*.
(**H**) UMAP plot of the cortical marker *FOXG1*.
(**I**) UMAP plot of the choroid plexus marker *TTR*.
(**J**) UMAP plot of the astrocyte marker *GFAP*. *GFAP* is also expressed in neural progenitors.
(**K**) UMAP plots of the upper-layer neuron markers *SATB2* and *CUX2*. *SATB2* is also expressed in subplate neurons.
(**L**) UMAP plot of the deep-layer neuron marker *BCL11B* (*CTIP2*).
(**M**) UMAP plot of the Layer IV marker *UNC5D*.
(**N**) UMAP plot of the Layer IV and upper Layer V marker *CRYM*.
(**O**) UMAP plot of the lower Layer V and Layer VI marker *FOXP2*.
For (**D-O**), color indicates the normalized transcript level.

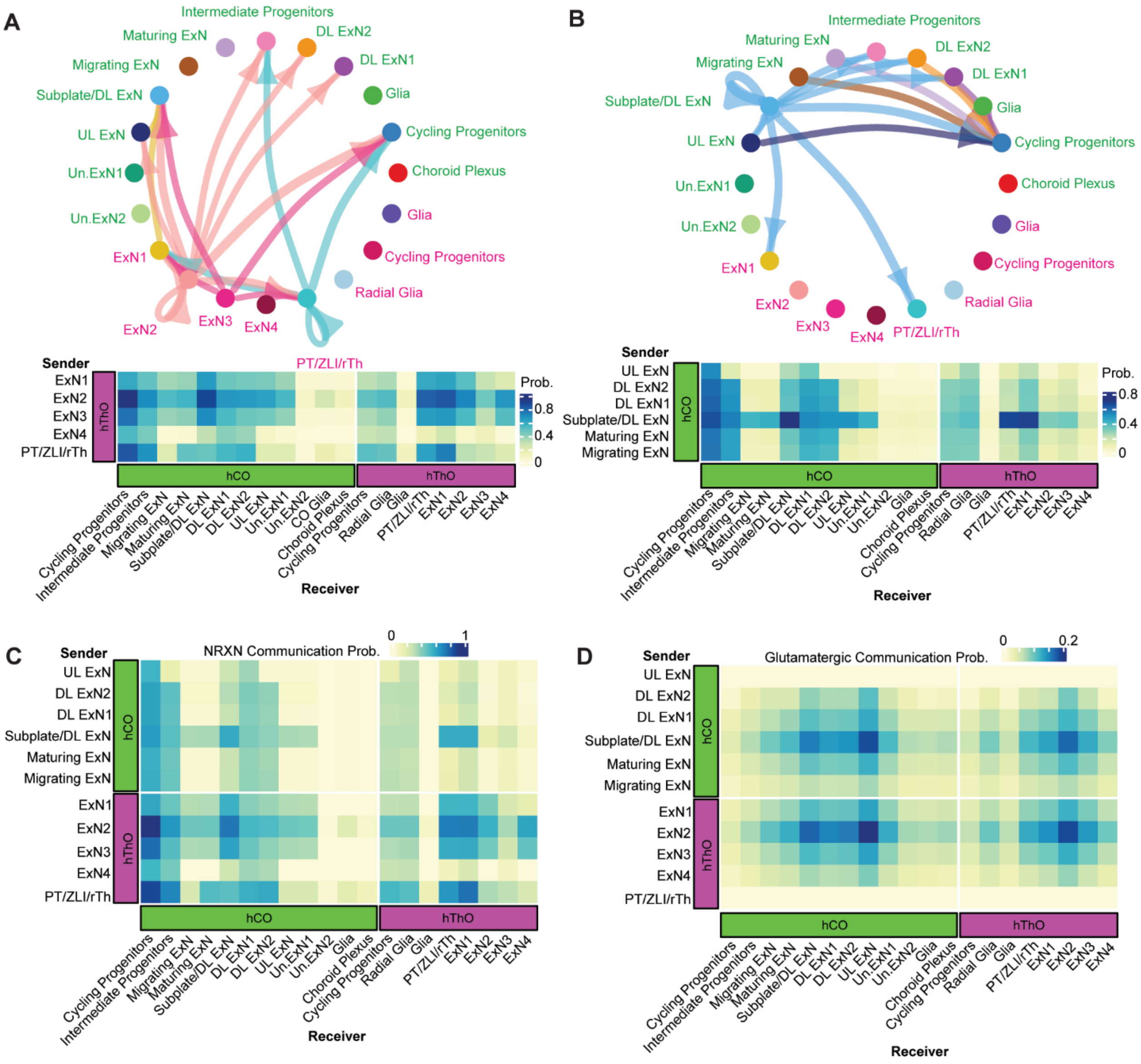


**Figure S6. NeuronChat predictions of communication between cell clusters in hThOs and hCOs (related to Figure 2).**

(**A**) NeuronChat predictions showing communication between cell clusters containing glutamatergic neurons in hThOs and other cell clusters in hCOs and hThOs. Probability of communication (Prob) is indicated in the heatmap (bottom). Interactions with a Prob >0.6 are shown in the circle plot (top).

(**B**) NeuronChat predictions showing communication between cell clusters containing glutamatergic neurons in hCOs and other cell clusters in hCOs and hThOs. Probability is indicated in the heatmap (bottom). Interactions with a Prob >0.5 are shown in the circle plot (top).

(**C**) Heatmap showing the probability of NRXN-mediated communication between cell clusters in hCOs and hThOs. Predictions were made using NeuronChat.

(**D**) Heatmap showing the probability of glutamatergic communication between cell clusters in hCOs and hThOs. Predictions were made using NeuronChat.

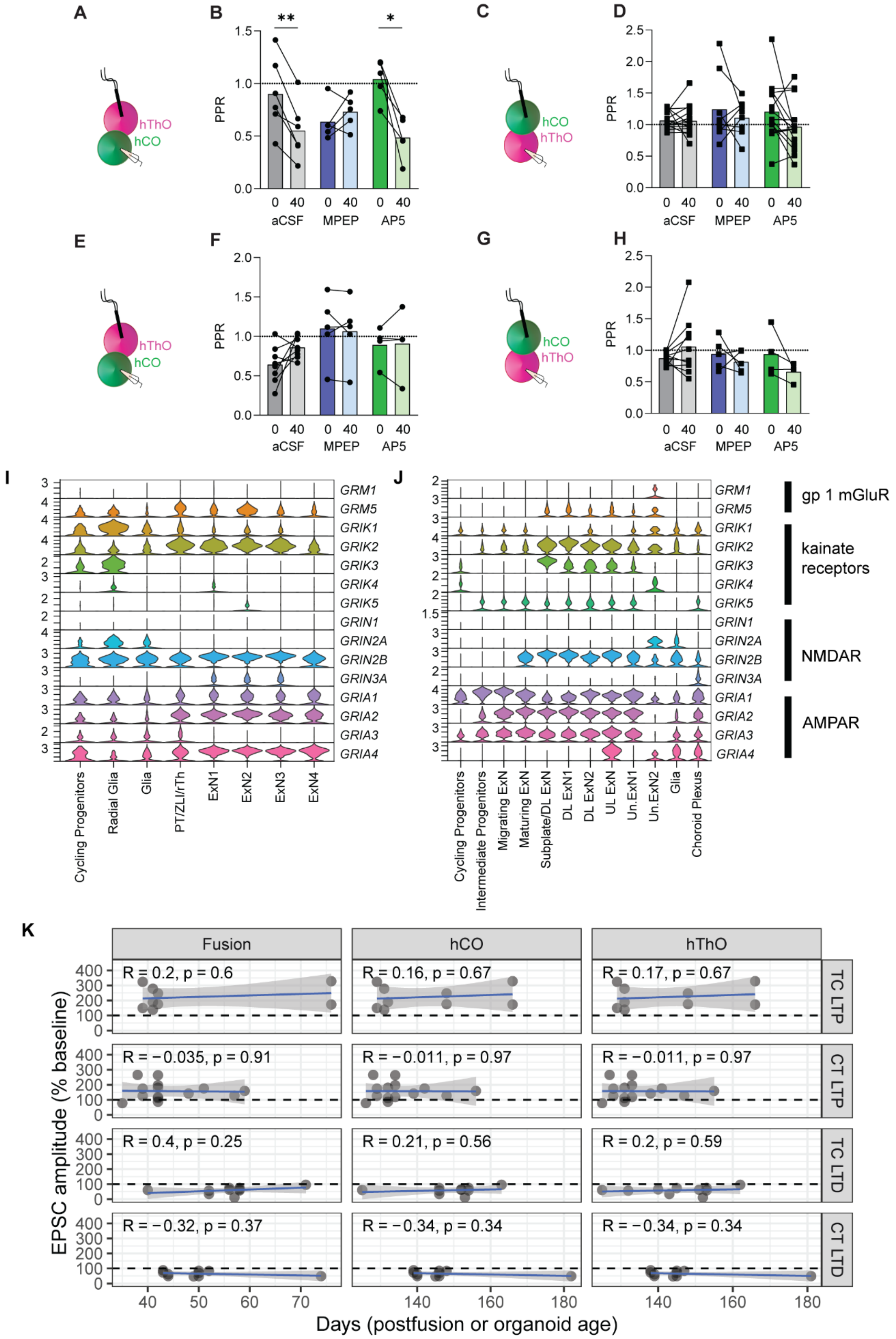

A
B
C
D
E
F
G
H
I
J
K
hThO
hCO
PPR
aCSF
MPEP
AP5
GRM1
GRM5
GRIK1
GRIK2
GRIK3
GRIK4
GRIK5
GRIN1
GRIN2A
GRIN2B
GRIN3A
GRIA1
GRIA2
GRIA3
GRIA4
gp 1 mGluR
kainate receptors
NMDAR
AMPAR
Cycling Progenitors
Radial Glia
Glia
PT/ZLI/rTh
ExN1
ExN2
ExN3
ExN4
Intermediate Progenitors
Migrating ExN
Maturing ExN
Subplate/DL ExN
DL ExN1
DL ExN2
UL ExN
Un.ExN1
Un.ExN2
Choroid Plexus
Fusion
hCO
hThO
TC LTP
CT LTP
TC LTD
CT LTD
R = 0.2, p = 0.6
R = 0.16, p = 0.67
R = 0.17, p = 0.67
R = −0.035, p = 0.91
R = −0.011, p = 0.97
R = −0.011, p = 0.97
R = 0.4, p = 0.25
R = 0.21, p = 0.56
R = 0.2, p = 0.59
R = −0.32, p = 0.37
R = −0.34, p = 0.34
R = −0.34, p = 0.34
EPSC amplitude (% baseline)
Days (postfusion or organoid age)

**Figure S7. Additional analyses related to TC and CT synaptic plasticities (related to Figures 3-7).**
(**A**) Schematic of the experimental condition for (**B**).
(**B**) Bar graph showing a change in the average PPR after TC LTP induction (35–40 min) compared to baseline (–5–0 min) in control aCSF (paired *t*-test, $^{**}p = 0.0028$). There is no change in PPR in 10 µM MPEP (paired *t*-test, $p = 0.34$), and there is a significant difference in PPR after LTP induction in 50 µM AP5 (paired *t*-test, $^{*}p = 0.01$).
(**C**) Schematic of the experimental condition for (**D**).
(**D**) Bar graph demonstrating that the average PPR values after CT LTP induction (40) are not different from baseline values (0) in control aCSF (paired *t*-test, $p = 0.97$), in 10 µM MPEP (paired *t*-test, $p = 0.49$), or in 50 µM AP5 (paired *t*-test, $p = 0.069$).
(**E**) Schematic of the experimental condition for (**F**).
(**F**) Bar graph showing that the average PPR after TC LTD induction (45) does not significantly differ from baseline PPR values (0) in control aCSF (paired *t*-test, $p = 0.082$), in 10 µM MPEP (paired *t*-test, $p = 0.66$), or in 50 µM AP5 (paired *t*-test, $p = 0.88$).
(**G**) Schematic of the recording configuration for (**H**).
(**H**) Bar graph demonstrating that the average PPR does not differ between baseline (0) and post–CT LTD induction (45) in control aCSF (paired *t*-test, $p = 0.188$), in 10 µM MPEP (paired *t*-test, $p = 0.477$), or in 50 µM AP5 (paired *t*-test, $p = 0.186$).
(**A-H**) Data are shown as the mean PPRs with individual values overlaid.
(**I**) Violin plot summarizing the levels of a subset of glutamate receptor transcripts by cell cluster in hThOs.
(**J**) Violin plot summarizing the levels of a subset of glutamate receptor transcripts by cell cluster in hCOs.
(**I** and **J**) Data were generated by snRNA-seq.
(**K**) Relationship between the average normalized post-induction EPSC amplitude and either the age of individual organoids or the post-fusion age (Fusion) of assembloids from control aCSF conditions. There is no significant effect of the individual organoid age or the age post-fusion on the expression of TC or CT LTP or LTD. R: Pearson correlation coefficient, p: p-value from Pearson correlation analysis.

### Supplemental Table

#### Table S1. CRISPR–Cas9 editing construct sequences (related to STAR Methods).

| Name | Sequence (5′ to 3 ′) |
|---|---|
| sgRNA Spacers | |
| TCF7L2-tdTomato | |
| RL14.TCF7L2.g10 | CCUCCACCGCCGUUCAGCUG |
| VGLUT1-tdTomato | |
| PC70.VGLUT1.g13 | UGGUCAGUAGUCCCGGACAG |
| Primers for PCR and Deep-Sequencing Validation | |
| TCF7L2-tdTomato | |
| RL14.hTCF7L2.5gen.F2 | TTCTAAAGTGCACTGTTTTGGGC |
| RL14.hTCF7L2.5junc.R2 | CCTCGCCCTCGATCTCGAAC |
| RL14.hTCF7L2.3junc.DS.F | CCCACAACGAGGACTACACC |

| | |
|---|---|
| RL14.hTCF7L2.3gen.R | AAAGCAAGAACGTGGCGAAG |
| RL14.DS.F2 | AAACTCACGCGTGCAGAAG |
| RL14.DS.R2 | GCCGAGGAGTTTTCGGAGC |
| VGLUT1-tdTomato | |
| PC70.hVGLUT1.5gen.F | TTCGCCATCTCTGGTGAGAAC |
| PC70.hVGLUT1.junc.DS.R | CACCTTGAAGCGCATGAACTC |
| PC70.hVGLUT1.junc.DS.F | CACCATCGTGGAACAGTACGA |
| PC70.hVGLUT1.3gen.R | GAGAGGGGAAGGATCCCAGAT |
| PC70.DS.F | TTCGTTGGCCATGACCAGCTGGCTG |
| PC70.DS.R | GCACTCTCCCTTGTTCCCAGTGCTGT |
| Donor sequences | |
| TCF7L2-tdTomato | |
| RL14_g10_tdTomato_P2A.dsDNA.donor | TTTCTGAAACCGCCCCCTCCCGGAGCAAGTCCCTGC<br>ACCCTCGCCCAGAATCCCGGGCTCGCACACACTCCG<br>CGCAGGCCGCTCCCCCTGCACACTCCTCCCTCCGTC<br>TCCCCCCGGCTTCCCCGCCCCTCTCTTCCTCCTTCTT<br>TCCCTCCTCCCTCTCCCGGCGCCCGAAAGGATCATT<br>GTTAGCCGCCCCCGCCCCGCCCACCCCGGCTGTTTA<br>TTTATGCACACGTCACTGGGCCGGCCCCGCCCTCCG<br>GCATCTCATTAAGGCAGTGTGTTCCTCTCGCCCTGTC<br>AATAATCTCCGCTCCCAGACTACTCCGTTCCTCCGG<br>ATTTCGATCCCCCTTTTTCTATCTGTCAATCAGCGCC<br>GCCTTTGAACTGAAAAGCTCTCAGTCTAACTTCAAC<br>TCACTCAAATCCGAGCGGCACGAGCACCTCCTGTAT<br>CTTCGGCTTCCCCCCCCCTTTGCTCTTTATATCTGAC<br>TTCTTGTTGTTGTTGGTGTTTTTTTTTTTTTTACCCCC<br>CTTTTTTATTTATTATTTTTTTGCACATTGATCGGATC<br>CTTGGGAACGAGAGAAAAAAGAAACCCAAACTCAC<br>GCGTGCAGAAGATCTCCCCCCCCTTCCCCTCCCCTC<br>CTCCCTCTTTTCCCCTCCCCAGGAGAAAAAGACCCC<br>CAAGCAGAAAAAAGTTCACCTTGGACTCGTCTTTTT<br>CTTGCAATATTTTTTGGGGGGGCAAAACTTTTTGGG<br>GGTGATTTTTTTTGGCTTTTCTTCCTCCTTCATTTTTC<br>TTCCAAAATTGCTGCTGGTGGGTGAAAAAAAAATG<br>GTGAGCAAGGGCGAGGAGGTCATCAAAGAGTTCAT<br>GCGCTTCAAGGTGCGCATGGAGGGCTCCATGAACG<br>GCCACGAGTTCGAGATCGAGGGCGAGGGCGAGGGC<br>CGCCCCTACGAGGGCACCCAGACCGCCAAGCTGAA<br>GGTGACCAAGGGCGGCCCCCTGCCCTTCGCCTGGGA<br>CATCCTGTCCCCCCAGTTCATGTACGGCTCCAAGGC<br>GTACGTGAAGCACCCCGCCGACATCCCCGATTACAA<br>GAAGCTGTCCTTCCCCGAGGGCTTCAAGTGGGAGCG<br>CGTGATGAACTTCGAGGACGGCGGTCTGGTGACCGT<br>GACCCAGGACTCCTCCCTGCAGGACGGCACGCTGAT<br>CTACAAGGTGAAGATGCGCGGCACCAACTTCCCCCC<br>CGACGGCCCCGTAATGCAGAAGAAGACCATGGGCT<br>GGGAGGCCTCCACCGAGCGCCTGTACCCCCGCGAC<br>GGCGTGCTGAAGGGCGAGATCCACCAGGCCCTGAA<br>GCTGAAGGACGGCGGCCACTACCTGGTGGAGTTCA |

| | |
|---|---|
| | AGACCATCTACATGGCCAAGAAGCCCGTGCAACTG<br>CCCGGCTACTACTACGTGGACACCAAGCTGGACATC<br>ACCTCCCACAACGAGGACTACACCATCGTGGAACA<br>GTACGAGCGCTCCGAGGGCCGCCACCACCTGTTCCT<br>GGGGCATGGCACCGGCAGCACCGGCAGCGGCAGCT<br>CCGGCACCGCCTCCTCCGAGGACAACAACATGGCC<br>GTCATCAAAGAGTTCATGCGCTTCAAGGTGCGCATG<br>GAGGGCTCCATGAACGGCCACGAGTTCGAGATCGA<br>GGGCGAGGGCGAGGGCCGCCCCTACGAGGGCACCC<br>AGACCGCCAAGCTGAAGGTGACCAAGGGCGGCCCC<br>CTGCCCTTCGCCTGGGACATCCTGTCCCCCCAGTTC<br>ATGTACGGCTCCAAGGCGTACGTGAAGCACCCCGCC<br>GACATCCCCGATTACAAGAAGCTGTCCTTCCCCGAG<br>GGCTTCAAGTGGGAGCGCGTGATGAACTTCGAGGA<br>CGGCGGTCTGGTGACCGTGACCCAGGACTCCTCCCT<br>GCAGGACGGCACGCTGATCTACAAGGTGAAGATGC<br>GCGGCACCAACTTCCCCCCCGACGGCCCCGTAATGC<br>AGAAGAAGACCATGGGCTGGGAGGCCTCCACCGAG<br>CGCCTGTACCCCCGCGACGGCGTGCTGAAGGGCGA<br>GATCCACCAGGCCCTGAAGCTGAAGGACGGCGGCC<br>ACTACCTGGTGGAGTTCAAGACCATCTACATGGCCA<br>AGAAGCCCGTGCAACTGCCCGGCTACTACTACGTGG<br>ACACCAAGCTGGACATCACCTCCCACAACGAGGAC<br>TACACCATCGTGGAACAGTACGAGCGCTCCGAGGG<br>CCGCCACCACCTGTTCCTGTACGGCATGGACGAGCT<br>GTACAAGGGAAGCGGAGCCACGAACTTCTCTCTGTT<br>AAAGCAAGCAGGAGACGTGGAAGAAAACCCCGGTC<br>CTATGCCGCAGCTCAATGGCGGTGGAGGGGATGAC<br>CTAGGCGCCAACGACGAACTGATTTCCTTCAAAGAC<br>GAGGGCGAACAGGAGGAGAAGAGCTCCGAAAACTC<br>CTCGGCAGAGAGGGATTTAGCTGATGTCAAATCGTC<br>TCTAGTCAATGAATCAGAAACGAATCAAAACAGCT<br>CCTCCGATTCCGAGGTAGGAAAAGCCCCTCGGGCTG<br>GTGGGGTTTTTTATCTGTTTCCTGGGCTTGGCAAATG<br>TTGCTGAAAGGGGAGAAATCGGGGCTGGGGGCGGC<br>GGCGGGGCCCGGCGGGCGGCGTGTGCGTACGGTGC<br>CACCATTGCAAAAACTTGTAACCCTGTTTTTTTCTAC<br>CCCCCCCTCGACCTCGCCGATTCTTTTTCTCCCCCTT<br>CTCCCCCTTCTGCGTGGCGTTTGCCCCTCGCCCTCCC<br>CACCTCCACCCCTCTGGGAAGGCGGAAAGACGGCC<br>TCCGCCTCGCTCCGAAAGTTTCCGAGACAAATCCCG<br>GGAAAGTTTGGAAGAAGGTGAGTACGCCCCGCGCG<br>CCCCGCAGCCGCCCGGAGCCGCCCCCCGGCCGGC<br>CGCCCCGCGCGCCCCGGCCCCGCGCCCGGCTCGGCC<br>TGGCCCTGCGCCGGCCCGGTCGGGGCGCCCGGCCCC<br>TCGGGGCACTTTCTAAAAAGTTTCTCCTCACTCTCTC<br>CCGCTCCGCGCGGCCGCCGCTGTCCCCTCGCCGCCC<br>CGCCATGTTAGCGGCCAAGAGGCAAGATGGAGGGC<br>TCTTTAAGGGGCCACCGTATCCCGGCTACCCCTTCA<br>TCATGATCCCCGACCTGACGAGC |
| VGLUT1-tdTomato | |

| | |
|---|---|
| PC70.g13_tdTomato_dsDNA.donor | CGCAGCTTGGGGGACTCTGGTCAGCTCTGTAGGGAA<br>CTTCCTGGCTCAGGAACGGGGCCTCTCAGCCTCACA<br>AAAGGTGGAACATGAGTGGCGGGGAGGGGAGAGG<br>GTTGAGAACGTCCCAGTGGGAATCACTCGAGTGAG<br>ATTCCGGTGTGACGCCTTCCGACTTGTCTCTCCGCCA<br>GGGTTCAACGTGAACCACCTGGACATAGCCCCGCGC<br>TACGCCAGCATCCTCATGGGCATCTCCAACGGCGTG<br>GGCACACTGTCGGGCATGGTGTGCCCCATCATCGTG<br>GGGGCCATGACTAAGCACAAGGTGGGTGCCGCAGT<br>CCTGACTCAGTCTGCCTCCAAGTGTGTGCTTCGTAC<br>CTAACTGACCACGTGGTGGCACTGTCTCTGGGGTGC<br>CATCCCTGCCTTTTCAGCCTCCCTGGGTCTCTGCCTC<br>CCTTCCTCTAGGCCTCCCCCTCTCTCCGCCTCTGTTT<br>CTCTGTCCTCCCATTTGACCTCAGCCCTATTGTCCCC<br>TGCAGACTCGGGAGGAGTGGCAGTACGTGTTCCTAA<br>TTGCCTCCCTGGTGCACTATGGAGGTGTCATCTTCTA<br>CGGGGTCTTTGCTTCTGGAGAGAAGCAGCCGTGGGC<br>AGAGCCTGAGGAGATGAGCGAGGAGAAGTGTGGCT<br>TCGTTGGCCATGACCAGCTGGCTGGCAGTGACGACA<br>GCGAAATGGAGGATGAGGCTGAGCCCCCGGGGGCA<br>CCCCCTGCACCCCCGCCCTCCTATGGGGCCACACAC<br>AGCACATTTCAGCCCCCCAGGCCCCCACCCCCTGTC<br>CGGGACTACGGAAGCGGAGCCACGAACTTCTCTCTG<br>TTAAAGCAAGCAGGAGACGTGGAAGAAAACCCCGG<br>TCCTGTGAGCAAGGGCGAGGAGGTCATCAAAGAGT<br>TCATGCGCTTCAAGGTGCGCATGGAGGGCTCCATGA<br>ACGGCCACGAGTTCGAGATCGAGGGCGAGGGCGAG<br>GGCCGCCCCTACGAGGGCACCCAGACCGCCAAGCT<br>GAAGGTGACCAAGGGCGGCCCCCTGCCCTTCGCCTG<br>GGACATCCTGTCCCCCCAGTTCATGTACGGCTCCAA<br>GGCGTACGTGAAGCACCCCGCCGACATCCCCGATTA<br>CAAGAAGCTGTCCTTCCCCGAGGGCTTCAAGTGGGA<br>GCGCGTGATGAACTTCGAGGACGGCGGTCTGGTGA<br>CCGTGACCCAGGACTCCTCCCTGCAGGACGGCACGC<br>TGATCTACAAGGTGAAGATGCGCGGCACCAACTTCC<br>CCCCCGACGGCCCCGTAATGCAGAAGAAGACCATG<br>GGCTGGGAGGCCTCCACCGAGCGCCTGTACCCCCGC<br>GACGGCGTGCTGAAGGGCGAGATCCACCAGGCCCT<br>GAAGCTGAAGGACGGCGGCCACTACCTGGTGGAGT<br>TCAAGACCATCTACATGGCCAAGAAGCCCGTGCAA<br>CTGCCCGGCTACTACTACGTGGACACCAAGCTGGAC<br>ATCACCTCCCACAACGAGGACTACACCATCGTGGAA<br>CAGTACGAGCGCTCCGAGGGCCGCCACCACCTGTTC<br>CTGGGGCATGGCACCGGCAGCACCGGCAGCGGCAG<br>CTCCGGCACCGCCTCCTCCGAGGACAACAACATGGC<br>CGTCATCAAAGAGTTCATGCGCTTCAAGGTGCGCAT<br>GGAGGGCTCCATGAACGGCCACGAGTTCGAGATCG<br>AGGGCGAGGGCGAGGGCCGCCCCTACGAGGGCACC<br>CAGACCGCCAAGCTGAAGGTGACCAAGGGCGGCCC<br>CCTGCCCTTCGCCTGGGACATCCTGTCCCCCCAGTTC<br>ATGTACGGCTCCAAGGCGTACGTGAAGCACCCCGCC |

| | |
|---|---|
| | GACATCCCCGATTACAAGAAGCTGTCCTTCCCCGAG<br>GGCTTCAAGTGGGAGCGCGTGATGAACTTCGAGGA<br>CGGCGGTCTGGTGACCGTGACCCAGGACTCCTCCCT<br>GCAGGACGGCACGCTGATCTACAAGGTGAAGATGC<br>GCGGCACCAACTTCCCCCCCGACGGCCCCGTAATGC<br>AGAAGAAGACCATGGGCTGGGAGGCCTCCACCGAG<br>CGCCTGTACCCCCGCGACGGCGTGCTGAAGGGCGA<br>GATCCACCAGGCCCTGAAGCTGAAGGACGGCGGCC<br>ACTACCTGGTGGAGTTCAAGACCATCTACATGGCCA<br>AGAAGCCCGTGCAACTGCCCGGCTACTACTACGTGG<br>ACACCAAGCTGGACATCACCTCCCACAACGAGGAC<br>TACACCATCGTGGAACAGTACGAGCGCTCCGAGGG<br>CCGCCACCACCTGTTCCTGTACGGCATGGACGAGCT<br>GTACAAGTGACCATGTGCCTCCCACTGAATGGCAGT<br>TTCCAGGACCTCCATTCCACTCATCTCTGGCCTGAGT<br>GACAGTGTCAAGGAACCCTGCTCCTCTCTGTCCTGC<br>CTCAGGCCTAAGAAGCACTCTCCCTTGTTCCCAGTG<br>CTGTCAAATCCTCTTTCCTTCCCAATTGCCTCTCAGG<br>GGTAGTGAAGCTGCAGACTGACAGTTTCAAGGATA<br>CCCAAATTCCCCTAAAGGTTCCCTCTCCACCCGTTCT<br>GCCTCAGTGGTTTCAAATCTCTCCTTTCAGGGCTTTA<br>TTTGAATGGACAGTTCGACCTCTTACTCTCTCTTGTG<br>GTTTTGAGGCACCCACACCCCCCGCTTTCCTTTATCT<br>CCAGGGACTCTCAGGCTAACCTTTGAGATCACTCAG<br>CTCCCATCTCCTTTCAGAAAAATTCAAGGTCCTCCTC<br>TAGAAGTTTCAAATCTCTCCCAACTCTGTTCTGCATC<br>TTCCAGATTGGTTTAACCAATTACTCGTCCCCGCCAT<br>TCCAGGGATTGATTCTCACCAGCGTTTCTGATGGAA<br>AATGGCGGTTTCAAGTCCCCGATTCCGTGCCCACTT<br>CACATCTCCCCTACCAGCAGATTCTGCGAAAGCACC<br>AAATTTCTCAAGACCCTCTTCTCCCTAGCTTAGCATA<br>ATGTCTGGGGAAACAACCAAAATCGCAATTTTAACA<br>ATATGCCTCTCTACCCCCGTGCACTTTTTCTGACATG<br>GTTTTCAGGTCTAAATAGTGGCTGCTCCAGTCCATG<br>AACTCAAAGGTTTGAAGCTACCACCATTGAACTCCC<br>CCATGGTG |

**Table S2. Number of independent batches of fused organoids used for synaptic plasticity experiments (related to Figures 4-7).**

| | TC LTP | CT LTP | TC LTD | CT LTD |
|---|---|---|---|---|
| **aCSF** | 3 | 2 | 4 | 2 |
| **AP5** | 2 | 2 | 3 | 2 |
| **MPEP** | 2 | 2 | 2 | 2 |
| **BAPTA** | 2 | 2 | 3 | 3 |
